# Targeting IL-6-STAT3/STAT4 Signaling Restores FOXP3 Expression in Pulmonary Arterial Endothelium and Reveals Novel Biomarkers for PAH

**DOI:** 10.64898/2026.04.29.721729

**Authors:** Prabhjot Kaur, Katherine Jankowski, Anurupa Ghosh, Carlos G Santos-Gallego, Anjira Ambade, Tahiris Duran, Yomna El Zibaoui, Maria T. Ochoa, Maryam Mansoori, Hooman Poor, Yassine Sassi, Paul M Hassoun, Malik Bisserier, Lahouaria Hadri

**Affiliations:** Department of Pharmacological Sciences, Icahn School of Medicine, Mount Sinai, New York, NY 10029, USA; Cardiology Department, CVRI, Icahn School of Medicine at Mount Sinai, New York, NY, USA; Division of Pulmonary and Critical Care Medicine, Johns Hopkins University, Baltimore, MD; Department of Cell Biology and Anatomy and Physiology, New York Medical College, Valhalla, NY, USA; Division of Pulmonology, Critical Care and Sleep Medicine, Icahn School of Medicine at Mount Sinai, New York, NY, USA; Fralin Biomedical Research Institute at Virginia Tech Carilion, Roanoke, Virginia, USA; Department of Biomedical Sciences and Pathobiology, Virginia-Maryland College of Veterinary Medicine, Virginia Tech, Blacksburg, Virginia; Department of Internal Medicine, VTC School of Medicine, Roanoke, Virginia

**Keywords:** Pulmonary hypertension, IL-6 neutralization, STAT3, FOXP3, DNA methylation

## Abstract

**Background:** Interleukin-6 (IL-6) is a central driver of pulmonary vascular remodeling in idiopathic, heritable, and connective tissue disease-associated pulmonary arterial hypertension (PAH). Elevated IL-6 correlates with right ventricular (RV) dysfunction and poor survival. However, the specific downstream mechanisms by which IL-6 drives pathogenesis remain poorly defined. We investigated the therapeutic impact of direct IL-6 neutralization and its regulation of a novel epigenetic signaling axis in PAH.

**Materials and Methods:** We evaluated species-specific IL-6-neutralizing antibodies in murine Sugen/hypoxia and rat monocrotaline models of PAH. RV structure and function were assessed using cardiac MRI and invasive hemodynamics. Lung transcriptomic profiling was performed by RNA sequencing mouse lung tissue. Key findings were validated in explanted human PAH lungs, serum, and peripheral blood mononuclear cells (PBMCs), and further interrogated through mechanistic in vitro studies in human pulmonary artery endothelial cells (PAECs).

**Results:** IL-6 neutralization significantly improved RV function, reduced pulmonary arterial pressures, and attenuated pulmonary vascular remodeling in both experimental models. Transcriptomic analysis identified a dysregulated FOXP3 signaling axis. Mechanistically, IL-6 induced cooperative binding of phosphorylated STAT3 and STAT4 to the *FOXP3* promoter, facilitating DNMT1-mediated DNA methylation and stable gene silencing. IL-6 blockade restored downstream FOXP3 expression, rescued downstream BMPR2 signaling, and re-established endothelial homeostasis. In clinical PAH cohorts, FOXP3 expression was markedly reduced and inversely correlated with circulating IL-6 levels and indices of disease severity.

**Conclusion:** IL-6 drives pulmonary hypertension through STAT3/STAT4- and DNMT1-dependent epigenetic repression of FOXP3, linking chronic inflammation to BMPR2 dysfunction and pulmonary vascular remodeling. IL-6 neutralization reverses this pathogenic program in experimental PAH. FOXP3 emerges as a mechanistic biomarker of disease severity and a potential tool for precision stratification of patients likely to benefit from IL-6-targeted therapies.

**Clinical Perspective:** *What Is New?:*

- **Synergistic Gene Repression:** The study demonstrates that phosphorylated STAT3 functional transcriptional complex that binds directly to the *FOXP3* promoter. Acting as a molecular scaffold, this complex actively represses FOXP3 expression within the pulmonary endothelium.
- **Epigenetic Silencing Mechanism:** IL-6 does not merely induce a transient suppression of FOXP3. Instead, IL-6/STAT3 signaling recruits DNMT1 to the *FOXP3* promoter, leading to site-specific DNA methylation and long-term epigenetic silencing. This mechanism provides a direct molecular link between chronic systemic inflammation and sustained pulmonary vascular injury.
- **The FOXP3-BMPR2 Connection:** The findings identify FOXP3 as a critical positive regulator of BMPR2 expression. Neutralization of IL-6 disrupts this pathological inhibitory loop, permitting restoration of BMPR2 signaling, the central gatekeeper of pulmonary vascular integrity and homeostasis.
- **Translational Biomarkers:** Importantly, the molecular changes within the lung are reflected systemically. Suppressed FOXP3 expression is readily detectable in PBMCs and serum, offering a non-invasive biomarker that correlates with pulmonary hemodynamic status and vascular health.

*Clinical Implications:*

- **Precision Patient Stratification:** FOXP3 expression in PBMCs and serum functions as a surrogate marker of pulmonary vascular epigenetic health. This could identify a specific inflammatory endotype of PAH patients most likely to respond to IL-6-targeted therapies.
- **Prognostic Biomarker:** Because FOXP3 levels integrate both systemic inflammatory burden and dysfunction of the RV-pulmonary vascular axis, they may provide a more sensitive tool for monitoring therapeutic response and predicting clinical worsening than current standard-of-care markers.

## INTRODUCTION

Pulmonary arterial hypertension (PAH) is a progressive disorder of the pulmonary vasculature characterized by elevated pulmonary arterial pressure and increased pulmonary vascular resistance, ultimately leading to right ventricular (RV) dysfunction and failure. If left untreated, the prognosis remains poor, with a median survival of approximately two to three years after diagnosis^1–3^. The disease is characterized by remodeling of distal pulmonary arteries, including endothelial dysfunction, proliferation of smooth muscle cells and fibroblasts, and accumulation of extracellular matrix ^1,4^. In parallel, the RV undergoes maladaptive remodeling, including cardiomyocyte hypertrophy and fibrosis, driven by sustained pressure overload and inflammatory signaling ^5,6^. Chronic inflammation is a key feature of PAH, with infiltration of inflammatory cells and increased circulating cytokines and chemokines, including interleukin-6 (IL-6)^7,8^. Current therapies mainly target vascular tone and endothelial function, providing symptomatic relief but failing to reverse vascular remodeling or significantly improve long-term survival, highlighting the urgent need for new therapeutic strategies ^9–12^. IL-6 is a key proinflammatory cytokine that is elevated in the serum and lung tissue of patients with PAH, with circulating levels correlating with disease severity and prognosis ^8,13,14^. Although IL-6 is known to be upregulated in pulmonary vascular cells in PAH, its precise cellular sources, target cells, and downstream signaling pathways remain incompletely defined^15–18^. Its established role in inflammatory and autoimmune diseases, including rheumatoid arthritis, chronic lung disease, and cancer, underscores its potential as a therapeutic target^19–26^.

Preclinical evidence from multiple experimental systems has consistently implicated IL-6 as a central mediator of pulmonary vascular inflammation, RV remodeling, and disease progression in pulmonary hypertension (PH) ^27–29^. Previous studies demonstrated that IL-6 expression is markedly elevated in the lungs, right ventricle, and circulating leukocytes in experimental PH models, including monocrotaline (MCT), hypoxia, and Sugen/hypoxia (SuHx), with levels correlating with disease severity and vascular remodeling ^29,30^. Genetic loss-of-function studies in IL-6 knockout mice revealed attenuated pulmonary vascular thickening, reduced inflammatory cell infiltration, and improved RV function, providing evidence that IL-6 drives PAH pathogenesis^11^. Therapeutic studies using IL-6-neutralizing antibodies or IL-6 receptor blockade across multiple species and models have repeatedly demonstrated that IL-6 inhibition suppresses STAT3 activation while attenuating PH ^12^. Together, these findings provided proof-of-concept that IL-6 is a viable therapeutic target in PAH. However, despite compelling preclinical evidence, the translational success of IL-6 blockade in PAH has been limited. The early-phase TRANSFORM-UK trial evaluated the IL-6 receptor-blocking monoclonal antibody tocilizumab in a small cohort of PAH patients^31^. While the therapy was well tolerated, it failed to achieve statistically significant improvements in key clinical endpoints, including pulmonary vascular resistance (PVR) and exercise capacity, in an unselected population ^32,33^. These findings highlight the heterogeneity of PAH pathobiology and suggest that global IL-6 inhibition may be insufficient without patient stratification by inflammatory phenotype or relevant molecular biomarkers Our work extends these observations by demonstrating that IL-6 neutralization robustly ameliorates pulmonary vascular remodeling and RV pathology across multiple preclinical PAH models. Importantly, we identify downstream effectors, most notably forkhead box P3 (FOXP3), whose regulation may refine therapeutic targeting. Integrating functional imaging, invasive hemodynamics, histology, and transcriptomics, we reveal that the IL-6-FOXP3 signaling axis is a critical mediator of pulmonary vascular remodeling. IL-6 blockade restores FOXP3 expression by relieving repression mediated by STAT3/STAT4 and DNA methylation, leading to improved RV function and reduced vascular pathology. Taken together, preclinical studies identify IL-6 as a key driver of PAH, while clinical data highlight the need for more targeted approaches. Our findings position FOXP3 as both a mechanistic effector and a potential biomarker to guide a precision medicine approach to IL-6-targeted therapies in PAH.

## MATERIALS AND METHODS

### Complete methods are in the Online Supplement

#### Patient and Sample Collection

Lung tissue specimens were obtained from patients enrolled in the Pulmonary Hypertension Breakthrough Initiative (PHBI); their characteristics are summarized in **Supplementary Table 1**. All samples were de-identified and archived, and their use did not constitute human subjects’ research. Tissue collection and analysis were conducted in accordance with protocols approved by the Institutional Review Boards at all participating PHBI sites. Peripheral blood samples and serum were collected from patients with PAH at Mount Sinai and Johns Hopkins University according to protocols approved by the Institutional Review Boards (**Supplementary Table 1 & 2**). Blood was processed to obtain serum using standard Ficoll-Paque density gradient centrifugation. Serum samples were aliquoted and stored at −80°C until analysis. PBMCs were either used immediately for functional assays or cryopreserved in liquid nitrogen for later experiments. Serum and peripheral blood mononuclear cells (PBMC) from healthy controls were obtained from ATCC and were either used immediately for functional assays or cryopreserved in liquid nitrogen for later experiments.

### Animals

All animal procedures were conducted in accordance with the National Institutes of Health (NIH) Guidelines for the Care and Use of Laboratory Animals and were approved by the Institutional Animal Care and Use Committee (IACUC) at the ISMMS.

### Sugen/Hypoxia (SuHx) Mouse Model of PH

PAH was induced in mice using the Sugen/hypoxia (SuHx) protocol, as previously described ^34,35^. Briefly, mice received subcutaneous injections of Semaxinib (SU5416; MedChem, Cat# HY-10374) at a dose of 20 mg/kg once weekly for three consecutive weeks. During this induction period, animals were maintained in hypoxic chambers with 10% oxygen. After three weeks, mice were returned to normoxic conditions (20% O₂) for the remainder of the study. Following the induction phase, SuHx mice were randomized into two treatment groups. The vehicle group (n = 7) received intraperitoneal (IP) injections of sterile-filtered PBS every three days for two weeks. The treatment group (n = 8) received IP injections of a 10 mg/kg monoclonal anti-mouse IL-6 neutralizing antibody (BE0046; InVivoMab, BioXCell) every 3 days for 2 weeks. A separate normoxic control group (n = 6) was maintained under standard oxygen conditions (20% O₂) for the duration of the experiment. At the end of the five-week study period, cardiac magnetic resonance imaging (cMRI) and hemodynamic measurements were obtained, and tissues were harvested for subsequent histological, molecular, and transcriptomic analyses.

### Monocrotaline (MCT) Rats Models of PH

Male Sprague Dawley rats were randomly assigned to a sham control group or an MCT group^34–38^. PH was induced in the MCT group by a single subcutaneous injection of monocrotaline (60 mg/kg; Sigma Aldrich). Three weeks after MCT administration, rats were randomized into two treatment groups. The anti-rIL-6 group (n = 7) received IP injections of an anti-rat IL-6 neutralizing antibody (AF506; R&D Systems) every 3 days for 2 weeks. The vehicle group (n = 7) received IP injections of sterile-filtered PBS on the same schedule. Following the treatment period, cMRI and hemodynamic assessments were performed, and tissues were collected for histologic, molecular, and transcriptomic analyses.

### Cardiac Magnetic Resonance Imaging and Hemodynamic Assessment

Cardiac structure and function were evaluated using both *in vivo* cMRI and invasive hemodynamic measurements, as previously described ^37^. For cMRI, rodents were imaged using a 7T Bruker small-animal scanner at the BioMedical Engineering and Imaging Institute (BME2I) at ISMMS. Anesthesia was maintained with 1.5-2% inhaled isoflurane via a nose cone, and body temperature was monitored throughout the session. Images were acquired using rodent-optimized sequences, including long-axis views of the right and left ventricles, short-axis stacks covering the entire ventricles for volumetric analysis, and four-chamber views for comprehensive assessment of all cardiac chambers. Dedicated software was used to quantify right and left ventricular end-diastolic and end-systolic volumes (EDV, ESV), stroke volume (SV), ejection fraction (EF), cardiac output (CO), ventricular wall thickness and mass, and structural remodeling.

Invasive hemodynamic measurements were performed following imaging. Animals were anesthetized with 2% isoflurane, placed on a heating pad to maintain normothermia, and intubated via tracheotomy. A surgical pericardial incision was made to expose the heart, and a flow probe (model 2.5S176, Transonic Systems, Ithaca, NY, USA) was positioned in the RV to measure RV systolic pressure (RVSP). In rats, mean pulmonary arterial pressure (mPAP) was assessed by advancing a catheter through the RV into the pulmonary artery, with correct placement confirmed by characteristic PA pressure waveforms^35–38^. Hemodynamic signals were recorded using a Scisense PV Control Unit (Transonic, Ontario, Canada), allowing continuous acquisition of RVSP, mPAP, and related parameters.

### Statistical Analysis

Data are presented as mean ± SEM. Data distribution was assessed for normality, and variance was evaluated for homogeneity. For comparisons between two groups, unpaired two-tailed Student’s T-tests were used. For multiple comparisons, one-way or two-way ANOVA followed by Tukey’s multiple comparisons test was applied as appropriate. Statistical analyses were performed using GraphPad Prism (GraphPad Software, Inc., La Jolla, CA). A P value <0.05 was considered statistically significant.

## RESULTS

### IL-6 neutralization improves RV function and alleviates pathological RV remodeling in the experimental MCT rat model of PH

IL-6 expression and signaling are elevated in PAH, implicating IL-6-mediated inflammation in disease progression ^11,12,27–29,39^. To assess the therapeutic potential of anti-rat IL-6 neutralizing antibodies (anti-rIL6 Ab), an MCT-treated rat model was used to induce an inflammatory response. In the MCT rat model, pulmonary vascular remodeling is characterized by hyperproliferation of pulmonary artery smooth muscle cells (PASMCs), endothelial dysfunction, medial thickening, and perivascular inflammation, processes in which IL-6 acts as a key pro-inflammatory and pro-proliferative mediator ^27,29^. After three weeks of MCT, animals received an anti-rIL6 or vehicle every three days for two weeks (**Figure 1A**). As expected, vehicle-treated MCT rats exhibited marked RV dysfunction, hypertrophy, and fibrosis, consistent with advanced PAH pathology (**Figure 1**). Treatment with anti-rIL6 Ab significantly improved RV structure and function compared to vehicle-treated MCT groups (**Figure 1B-C & Supplemental Table 1**). At study endpoints, cMRI showed improved RV systolic function in anti-rIL-6 Ab–treated rats, with increased RV longitudinal strain (RV-LS; 18.01 ± 1.55% vs. 14.41 ± 2.01%), higher ejection fraction (RVEF; 50.48 ± 5.46% vs. 36.79 ± 9.39%), greater tricuspid annular plane systolic excursion (TAPSE; 2.98 ± 0.37 mm vs. 2.18 ± 0.37 mm), and increased RV/PA coupling (1.02 ± 0.23 vs. 0.61 ± 0.20) (**Figure 1B-C & Supplemental Table 3**). Furthermore, anti-rIL-6 blockade reduced RV wall thickness (0.69 ± 0.17 mm vs. 0.98 ± 0.19 mm), decreased RV end-systolic volume (0.28 ± 0.054 mL vs. 0.38 ± 0.058 mL), reduced RV/LV ratio (1.24 ± 0.4 vs. 1.87 ± 0.44), and estimated pulmonary vascular resistance (PVR; 4.65 ± 067 WU vs. 6.69 ± 0.96 WU) compared to vehicle-treated MCT controls (**Figure 1B-C & Supplemental Table 3**). Notably, left ventricular (LV) function and dimensions were unchanged between groups **(Supplemental Table 4)**, indicating that IL-6 neutralization specifically ameliorates RV function in this model.

**Figure 1.**
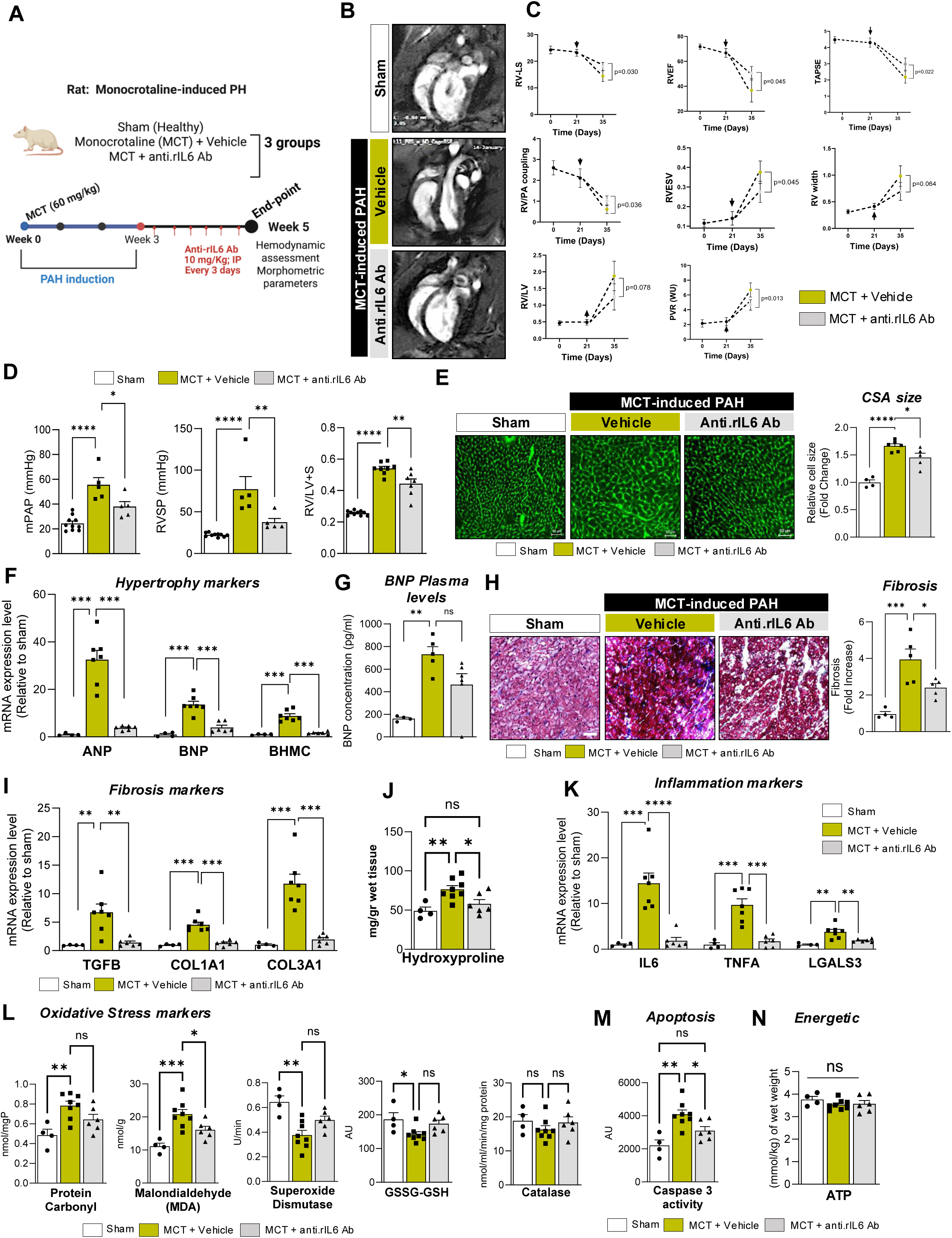
IL-6 blockade attenuates monocrotaline (MCT)-induced pulmonary arterial hypertension (PAH). (**A**) Experimental schematic illustrating the therapeutic protocol. Rats received a single monocrotaline (MCT) injection to induce PAH. After 3 weeks, animals were treated with a rat-specific IL-6 neutralizing antibody (anti-rIL-6 Ab; 20 mg/kg, intraperitoneally, every other day) for an additional 2 weeks. Tissues were collected at week 5 for molecular, biochemical, and histological analyses. (**B**) Representative long-axis cardiac magnetic resonance imaging (cMRI) images from the indicated groups following anti-rIL-6 antibody treatment. (**C**) Quantitative cMRI-derived cardiac hemodynamic parameters in sham controls (n = 4), MCT-PAH rats treated with vehicle (n = 4), or anti-rIL6 Ab (n = 4). RV-LS indicates right ventricular longitudinal strain; RVEF, RV ejection fraction; TAPSE, tricuspid annular plane systolic excursion; RV/PA coupling, right ventricular-pulmonary arterial coupling; RVESV, RV end-systolic volume; RV width indicates RV wall thickness; RV/LV ratio, right ventricular/left ventricular size ratio; Estimated PVR (WU), pulmonary vascular resistance in Wood units. (**D**) Mean pulmonary arterial pressure (mPAP), right ventricular systolic pressure (RVSP), and Fulton index [RV/(LV + S)] were measured in sham rats and MCT-treated rats receiving vehicle or anti-rIL-6 Ab. (**E**) Representative right ventricular (RV) sections stained with fluorescence-conjugated wheat germ agglutinin to assess cardiomyocyte cross-sectional area (left). Quantification of cardiomyocyte size is shown (n = 4-6). Scale bar = 50 μm. (**F**) Relative expression of cardiac hypertrophy–associated genes (ANP, BNP, β-MHC) in the indicated groups determined by RT-qPCR (n = 4-7). (**G**) Plasma BNP concentrations measured in the indicated experimental groups (n = 4–6). (**H**) Representative Masson’s trichrome–stained RV sections demonstrating myocardial fibrosis (left), with corresponding quantification (n = 4–5). Scale bar = 100 μm. (**I**) Expression of fibrosis-related genes (TGFB, COL1A1, COL3A1) assessed by RT-qPCR in RV tissue (n = 4-7). (**J**) RV hydroxyproline content as a measure of total collagen deposition (n = 4-7). (**K**) Relative expression of inflammatory markers (IL-6, TNFα, and galectin-3 [LGALS3]) determined by qPCR in RV tissue (n = 4-7). (**L**) Levels of oxidative stress–related markers (protein carbonyls, malondialdehyde [MDA], superoxide dismutase [SOD], GSSG/GSH ratio, and catalase activity) were measured in whole-lung homogenates. (**M-N**) Caspase-3 activity and ATP levels were measured in whole-lung homogenates of the indicated groups. Data are presented as mean ± SEM. *P < 0.05, **P < 0.01, ***P < 0.001.

### IL-6 neutralization reverses established PAH and attenuates RV remodeling in the MCT rat model

Elevated RVSP and RV hypertrophy are hallmark features of experimental and clinical PAH and reflect the hemodynamic burden imposed by increased pulmonary vascular resistance ^40,41^. Right heart catheterization revealed that anti-rIL6 Ab treatment markedly decreased mPAP and RVSP compared with vehicle-treated MCT rats (**Figure 1D**, left and middle panel), indicating improved cardiopulmonary function. RV hypertrophy, assessed by the Fulton Index, was also significantly reduced, reflecting improved RV adaptation to decreased afterload (**Figure 1D**, right panel). Histological analysis demonstrated a reduction in RV cardiomyocyte cross-sectional area in anti-rIL6 Ab-treated animals (**Figure 1E)**. Molecular markers of hypertrophy, including atrial natriuretic peptide (ANP), brain natriuretic peptide (BNP), and β-myosin heavy chain (β-MHC), were significantly lower in treated animals (**Figure 1F**). Notably, plasma BNP levels were also reduced, confirming attenuation of maladaptive RV stress responses (**Figure 1G**). Consistent with these findings, IL-6 blockade significantly reduced RV fibrosis, accompanied by decreased expression of pro-fibrotic genes, including transforming growth factor-β1 (*Tgfb1*), collagen type I alpha 1 (*Col1a1*), and collagen type III alpha 1 (*Col3a1*), in anti-rIL-6 Ab-treated MCT rats (**Figure 1H-I**). Moreover, collagen content, measured by hydroxyproline assay, was reduced, along with pro-inflammatory cytokines, including *Il6*, tumor necrosis factor-α (*Tnfa*), and galectin-3 (*Lgals3*) (**Figure 1J-K**). Furthermore, anti-rIL6 Ab treatment decreased levels of proteins associated with oxidative stress and cell death, indicating suppression of RV damage without changes in energy supply (ATP/ADP) (**Figure 1L-N**). Collectively, these results show that IL-6 neutralization reverses established features of PAH, attenuates RV hypertrophy and fibrosis, suppresses inflammatory signaling, and limits oxidative stress, supporting a central role for IL-6 in maladaptive RV remodeling in the MCT rat model.

### IL-6 Blockade Reduces Pulmonary Muscularization and Vascular Remodeling in MCT-induced PH

Anti-rIL6 Ab treatment attenuated medial thickening of distal pulmonary arteries and prevented lumen narrowing compared with vehicle-treated MCT rats (**Figure 2A**). Compared to the sham group, α-Smooth muscle actin (α-SMA, green) immunostaining showed robust expression throughout the vascular wall in vehicle-treated MCT rats, consistent with enhanced PASMC proliferation and distal arterial muscularization **(Figure 2B)**. In contrast, anti-rIL6 Ab-treated animals exhibited reduced medial thickness, decreased α-SMA staining intensity, and increased apoptotic markers, indicating suppression of SMC proliferation and vascular remodeling (**Figure 2B, Supplementary Figure 1A, respectively**). Together, these findings show that IL-6 neutralization reduces medial thickening and muscularization of distal pulmonary arteries, supporting its therapeutic potential in limiting vascular remodeling in experimental PH.

**Figure 2.**
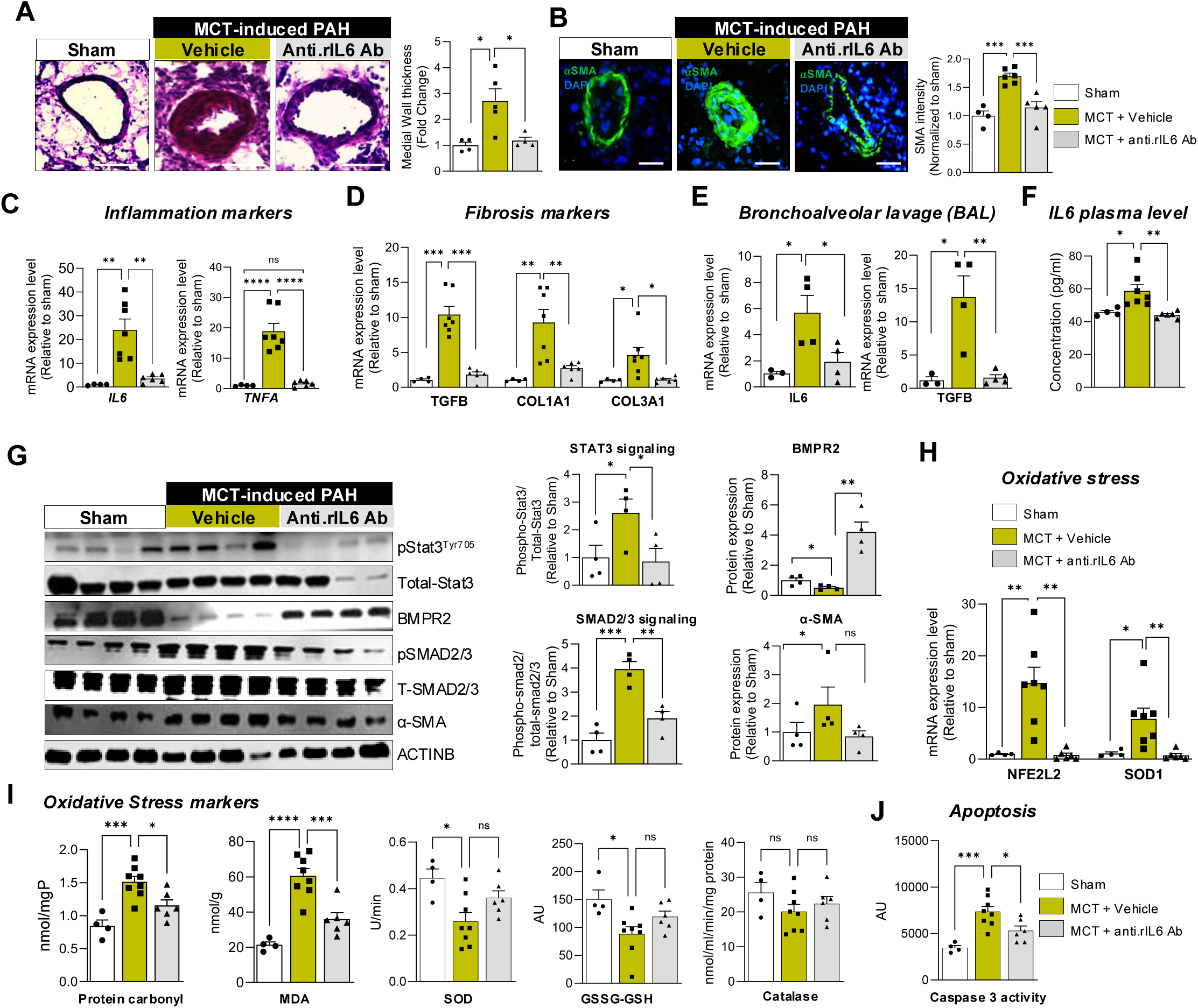
Neutralization of IL-6 attenuates pulmonary vascular remodeling in monocrotaline (MCT)-induced PAH. (**A**) Representative hematoxylin and eosin–stained lung sections from saline-treated control rats and MCT-challenged rats treated with vehicle or anti-rat IL-6 neutralizing antibody (anti-rIL-6 Ab) (left). Quantification of medial wall thickness of distal pulmonary arteries is shown (right). Five vessels were analyzed per animal, with four to six animals per group. (**B**) Representative immunofluorescence staining for α–smooth muscle actin (α-SMA, green) in lung sections from the indicated groups. Nuclei are counterstained with DAPI (blue). Scale bar = 100 μm. Quantification of distal pulmonary vascular α-SMA content is shown in the right panel (n = 4-7). (**C**) Relative expression of inflammatory genes (IL-6, TNFα) in lung tissue assessed by RT-qPCR in the indicated groups (n = 4-7). (**D**) Relative expression of fibrosis-associated genes (TGFβ, COL1A1, COL3A1) in lung tissue assessed by RT-qPCR (n = 4–7). (**E**) IL-6 and TGFβ mRNA expression levels measured in bronchoalveolar lavage (BAL) samples by RT-qPCR in the indicated groups (n = 3-4). (**F**) Plasma IL-6 concentrations measured by ELISA in the indicated groups (n = 4-7). (**G**) Representative immunoblots showing phosphorylated STAT3 (p-STAT3-T705), total STAT3, BMPR2, phosphorylated SMAD2/3 (p-SMAD2/3), total SMAD2/3, α-SMA, and β-actin (ACTINB) in lung homogenates (n = 4). Corresponding densitometric quantification is shown (right), with protein levels normalized to ACTINB. Phosphorylated STAT3 and SMAD2/3 levels were normalized to their respective total protein levels. (**H**) Relative lung mRNA expression levels of oxidative stress–related genes NFE2L2 (NRF2) and SOD1 assessed by RT-qPCR (n = 4-7). (**I**) Levels of oxidative stress–associated markers, including protein carbonyls, malondialdehyde (MDA), SOD1 activity, GSSG/GSH ratio, and catalase activity, measured in whole-lung homogenates (n = 4-8). (**J**) Caspase-3 activity measured in whole-lung homogenates of the indicated groups (n = 4-8). Data are presented as mean ± SEM. *P < 0.05, **P < 0.01, ***P < 0.001.

### IL-6 Neutralization Suppresses Pulmonary Inflammation, Fibrosis, and Pro-Proliferative Signaling in MCT-Induced PH

IL-6, a central mediator in PAH pathogenesis, activates STAT3 and interacts with TGF-β/SMAD2/3 signaling, promoting PASMC proliferation and extracellular matrix production ^12,27–29,42^. To assess the effects of IL-6 neutralization, lung tissues, serum, and bronchoalveolar lavage (BAL) fluid were analyzed in sham, vehicle- and anti-rIL6 Ab-treated MCT rats. Lung tissue from vehicle-treated MCT rats exhibited robust upregulation of pro-inflammatory transcripts, including IL-6 and TNF-α, and pro-fibrotic markers, such as *Tgfβ1*, *Col1a1*, and *Col3a1*, relative to sham controls (**Figure 2C-D**). Anti-rIL6 Ab treatment significantly reduced the expression of these markers, consistent with effective suppression of lung inflammation and fibrosis (**Figure 2C-D**). Similarly, elevated *Il6* and *Tgfβ1* mRNA levels in BAL fluid, as well as IL-6 plasma levels, were consistently decreased following IL-6 neutralization (**Figure 2E-F**). At the protein level, anti-rIL-6 Ab treatment reduced phosphorylation of signal transducer and activator of transcription 3 (STAT3) and SMAD family members 2 and 3 (SMAD2/3). α-SMA expression in lung homogenates also decreased, while BMPR2 protein levels were markedly increased compared to vehicle-treated animals, exceeding levels observed in sham controls (**Figure 2G**). These changes indicate reduced pro-proliferative and pro-fibrotic signaling and partial rebalancing of BMPR2/TGF-β pathways.

Immunohistochemical analysis of CD68 showed reduced macrophage infiltration in perivascular regions following IL-6 neutralization (**Supplementary Figure 1B**). Consistent with this, cytokine profiling using a Proteome Profiler cytokine array showed reduced signal intensity for CXCL7 and ICAM-1 in anti-rIL-6 Ab-treated lungs (**Supplementary Figure 2A-B**). IL-6 blockade also decreased mRNA and protein markers associated with response to oxidative stress, cellular injury, and apoptosis (**Figure 2I-J**), consistent with reduced tissue damage. Together, these findings show that IL-6 neutralization suppresses pulmonary inflammation, fibrosis, and vascular proliferation, supporting a central role for IL-6 in pulmonary vascular remodeling in MCT-induced PH.

### IL-6 neutralization improves RV function and attenuates structural remodeling in the SuHx mouse model of PH

To validate the therapeutic effects of IL-6 neutralization, SuHx mice used as a second PAH model, were treated every 3 days with a mouse-specific anti-IL-6 antibody for two weeks (**Figure 3A**). Cardiac MRI revealed marked structural and functional abnormalities in vehicle-treated SuHx mice compared with normoxic controls (**Figure 3B-C, Supplementary Table 5-6**). Representative long-axis ventricular images demonstrate these alterations across treatment groups (**Figure 3B**).

**Figure 3.**
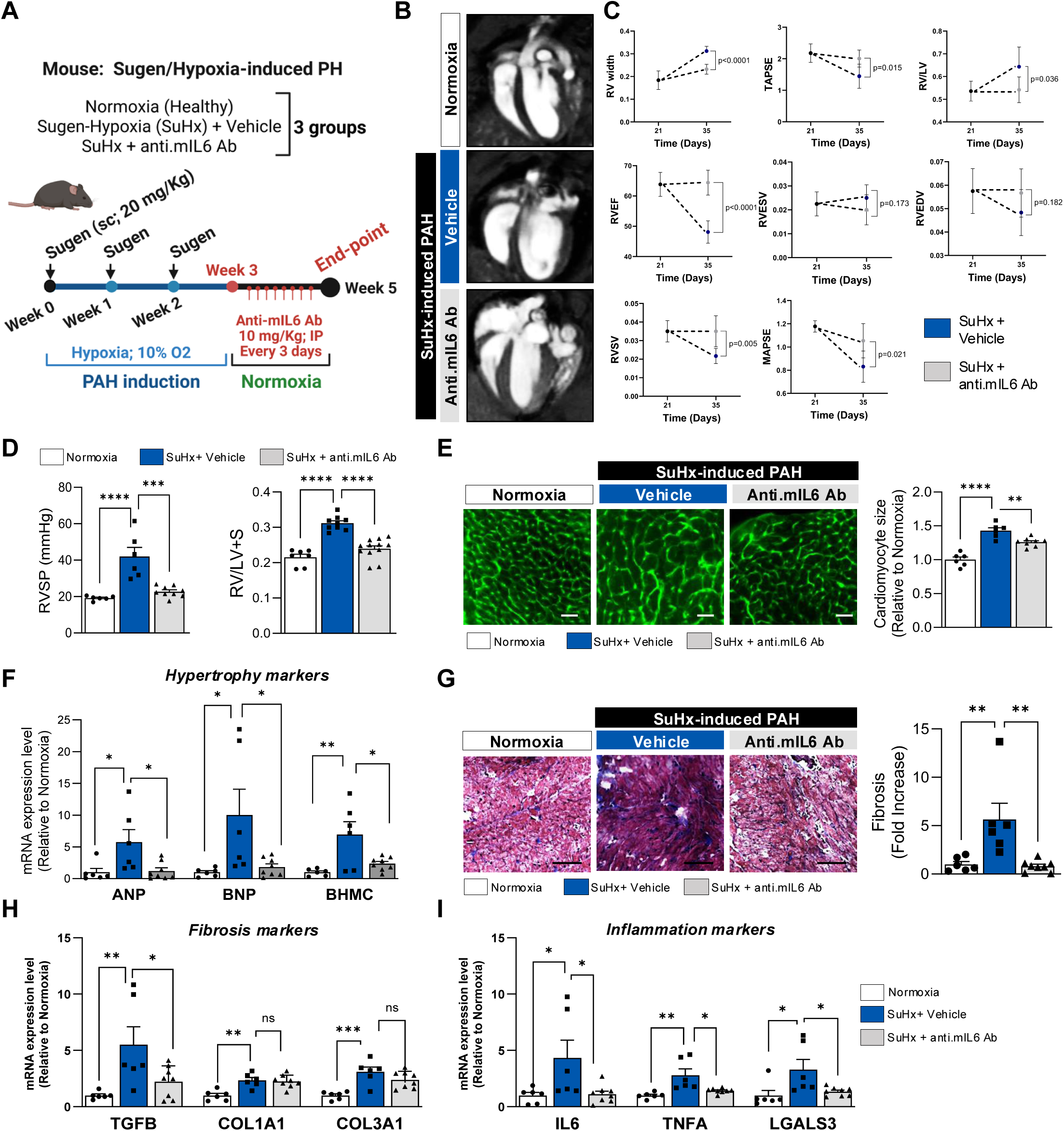
Neutralization of IL-6 attenuates right ventricular (RV) dysfunction and remodeling in the Sugen/hypoxia (SuHx)–induced PAH mouse model. **(A)** Schematic of experimental design. Mice were exposed to SuHx for 3 weeks, followed by 1 week of normoxia. Beginning thereafter, animals received a mouse-specific IL-6 neutralizing monoclonal antibody (anti-mIL6 Ab; 10 mg/kg, intraperitoneally, every other day) or vehicle for 2 weeks. Tissues were collected at week 6 for molecular and histological analyses. **(B)** Representative long-axis cardiac magnetic resonance imaging (cMRI) images from the indicated experimental groups following anti-mIL6 antibody treatment. **(C)** cMRI-derived cardiac hemodynamic parameters measured longitudinally in normoxic control mice (Nx, n = 4) and SuHx-PAH mice treated with vehicle (n = 6) or anti-mIL6 Ab (n = 6), including baseline, post-SuHx exposure, pre-treatment, and at the 5-week endpoint. RV width indicates right ventricular wall thickness; TAPSE, tricuspid annular plane systolic excursion; RV/LV ratio, right ventricular/left ventricular size ratio; RVEF, RV ejection fraction; RVESV, RV end-systolic volume; RVEDV, right ventricular end-diastolic volume; RVSV, right ventricular stroke volume; MAPSE, mitral annular plane systolic excursion. **(D)** Right ventricular systolic pressure (RVSP) and Fulton index [RV/(LV + S)] measured in normoxic control mice and SuHx-induced PAH mice treated with vehicle or anti-mIL6 Ab (n = 6-9). **(E)** Representative RV sections stained with fluorescence-conjugated wheat germ agglutinin (WGA) to assess cardiomyocyte cross-sectional area (left). Quantification of cardiomyocyte size is shown (n = 6-8). Scale bar = 50 μm. **(F)** Relative expression of cardiac hypertrophy–associated genes (ANP, BNP, β-MHC) in RV tissue assessed by RT-qPCR in the indicated groups (n = 6-8). **(G)** Representative Masson’s trichrome–stained RV sections illustrating myocardial fibrosis (left) with corresponding quantification (right) (n = 6-8). Scale bar = 100 μm. **(H)** Relative expression of fibrosis-related genes (TGFβ, COL1A1, COL3A1) in RV tissue assessed by RT-qPCR (n = 6-8). **(I)** Relative expression of inflammatory genes (IL-6, TNFα, and galectin-3 [LGALS3]) measured by RT-qPCR in lung tissue from normoxic and SuHx-induced PAH mice treated with vehicle or anti-mIL6 Ab (n = 6-8). Data are presented as mean ± SEM. *P < 0.05, **P < 0.01, ***P < 0.001.

Quantitative cMRI analysis showed increased RV width in the vehicle-treated SuHx mice at the 5-week endpoint (0.31 ± 0.02 g vs. 0.18 ± 0.04 g in controls), which was reduced by anti-mIL6 Ab treatment (0.23 ± 0.02 g), indicating attenuation of RV hypertrophy (**Figure 3C**). IL-6 blockade also restored TAPSE (2.005 ± 0.27 mm vs. 1.44 ± 0.38 mm), RVEF (64.41 ± 4.0% vs. 48.11 ± 3.68%), RVESV (0.035 ± 0.008 mL vs. 0.02 ± 0.004 mL), RVEDV (0.05 ± 0.01 mL vs. 0.04 ± 0.009 mL), and MAPSE (1.05 ± 0.15 mm vs. 0.83 ± 0.13 mm), while decreasing RVSV (0.035 ± 0.008 mL vs. 0.021 ± 0.04 mL) and RV/LV (0.54 ± 0.05 vs. 0.64 ± 0.08) compared with vehicle-treated SuHx mice **(Figure 3C, Supplementary Table 5),** indicating improvement in RV function, hypertrophy and RV remodeling. Left ventricular structure and function were unchanged across groups (**Supplementary Table 6**), indicating that IL-6 neutralization selectively improves RV remodeling in the SuHx model.

### IL-6 neutralization attenuates RV hypertrophy, fibrosis, and inflammation in SuHx mice

Hemodynamic measurements showed that RVSP was increased in vehicle-treated SuHx mice compared with normoxic controls and was reduced by anti–mIL-6 Ab treatment (**Figure 3D**). The Fulton index (RV/[LV+S]) was also decreased with IL-6 neutralization, indicating reduced RV hypertrophy (**Figure 3D**, left panel). Consistently, RV cardiomyocyte cross-sectional area was lower in anti-mIL6 Ab-treated mice than in vehicle-treated SuHx animals (**Figure 3E**). Markers of hypertrophy, including *Anp*, *Bnp*, and *βmhc*, were increased in vehicle-treated SuHx mice and normalized following IL-6 blockade (**Figure 3F**). Masson’s Trichrome staining showed extensive interstitial fibrosis in the RV of SuHx-vehicle mice, which was substantially reduced by the anti-mIL6 Ab-treatment (**Figure 3G**). Accordingly, pro-fibrotic genes, including Tgfb1, Col1a1, and Col3a1, decreased in the RV after IL-6 neutralization (**Figure 3H**). Pro-inflammatory markers, including *Tnfa, Il6,* and *Lgals3*, were elevated in SuHx-vehicle mice and reduced with anti–mIL-6 Ab treatment (**Figure 3I**). Together, these findings show that IL-6 blockade reduces RV hypertrophy, fibrosis, and inflammation, supporting its role in limiting maladaptive RV remodeling in SuHx-induced PAH.

### IL-6 neutralization reverses pulmonary vascular remodeling in SuHx mice

To assess the effect of IL-6 blockade on pulmonary vascular remodeling, lung sections from SuHx mice were analyzed by histology and immunohistochemistry. Anti-mIL6 Ab treatment reduced medial hypertrophy and wall thickness compared with vehicle-treated SuHx mice (**Figure 4A**). α-SMA immunostaining showed marked distal pulmonary artery muscularization in the SuHx-vehicle group, which was attenuated by anti-mIL6 treatment, along with increased medial cell apoptosis (**Figure 4B & Supplementary Figure 3A**).

**Figure 4.**
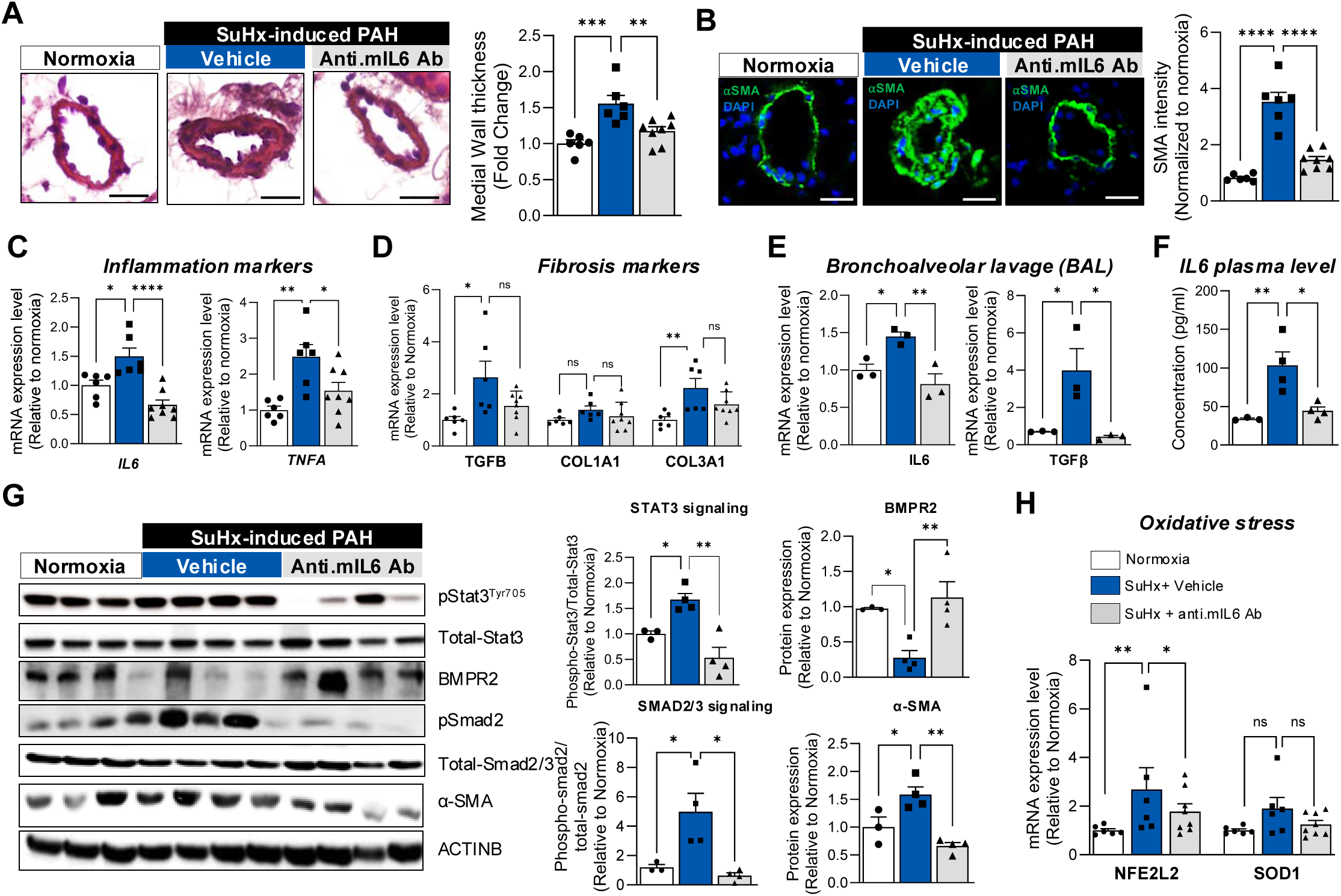
Post-treatment with a mouse IL-6 neutralizing antibody improves pulmonary vascular remodeling in the Sugen/hypoxia (SuHx)–induced PAH mouse model. **(A)** Representative hematoxylin and eosin (H&E)-stained lung sections from normoxic control mice and SuHx-challenged mice treated with vehicle or anti–mouse IL-6 neutralizing antibody (anti-mIL6 Ab) (left). Quantification of medial wall thickness of distal pulmonary arteries is shown (right). Five vessels were analyzed per animal, with six to eight animals per group. **(B)** Representative immunofluorescence staining for α–smooth muscle actin (α-SMA, green) in lung sections from the indicated groups. Nuclei were counterstained with DAPI (blue). Scale bar = 100 μm. Quantification of distal pulmonary vascular α-SMA content is shown (right). Five vessels were analyzed per animal, with six to eight animals per group. **(C)** Relative lung mRNA expression levels of inflammatory markers (IL-6, TNFα) assessed by RT-qPCR in the indicated groups (n = 6-8). **(D)** Relative lung mRNA expression levels of fibrosis-associated genes (TGFβ, COL1A1, COL3A1) assessed by RT-qPCR (n = 6-8). **(E)** IL-6 and TGFβ mRNA expression levels measured in bronchoalveolar lavage (BAL) samples by RT-qPCR (n = 3). **(F)** Plasma IL-6 concentrations measured by mouse-specific ELISA in the indicated groups (n = 3-4). **(G)** Representative immunoblots showing phosphorylated STAT3 (p-STAT3^T705), total STAT3, BMPR2, phosphorylated SMAD2/3 (p-SMAD2/3), total SMAD2/3, α-SMA, and β-actin (ACTINB) in lung homogenates. Corresponding densitometric quantification is shown (right). Protein levels were normalized to ACTINB, and phosphorylated proteins were normalized to their respective total protein levels (n = 3-4). **(H)** Relative lung mRNA expression levels of oxidative stress–related genes NFE2L2 (NRF2) and SOD1 assessed by RT-qPCR in the indicated groups (n = 6-8). Data are presented as mean ± SEM. *P < 0.05, **P < 0.01, ***P < 0.001.

Molecular profiling of lung tissue showed increased expression of pro-inflammatory and pro-fibrotic markers, including *Il6, Tnf, Tgfb1, Col1a1,* and *Col3a1*, in SuHx-vehicle mice, which was reduced by IL-6 neutralization (**Figure 4C-D**). Similarly, *Il6* and *Tgfb1* mRNA levels in BAL fluid and circulating IL-6 in plasma were decreased following IL-6 blockade (**Figure 4E-F**). At the signaling level, phosphorylation of STAT3 and SMAD2/3 and α-SMA expression were increased in SuHx lungs and reduced by anti-mIL-6 Ab treatment, while BMPR2 levels were restored, indicating inhibition of pro-proliferative and pro-fibrotic pathways (**Figure 4G**). Immunohistochemical analysis revealed increased perivascular macrophage accumulation (CD68) in SuHx-vehicle mice, which was reduced following IL-6 neutralization, along with increased apoptosis (**Supplementary Figure 3A-B**). Cytokine profiling using a Proteome Profiler mouse cytokine array showed reduced signal intensity for CXCL7 and ICAM-1, along with decreased markers of oxidative damage, in anti-mIL-6 Ab-treated lungs compared with vehicle-treated SuHx mice (**Supplementary Figure 4A-B, Figure 4H**, respectively). Collectively, these findings demonstrate that IL-6 neutralization exerts broad molecular and cellular effects in the SuHx lung by suppressing inflammation, reducing fibrotic signaling, inhibiting PASMC proliferation, and limiting immune cell infiltration, thereby attenuating pulmonary vascular remodeling.

### RNA sequencing identifies FOXP3 as a key transcriptional target repressed by IL-6 in PH

To define the global transcriptional consequences of IL-6 neutralization in PAH, we performed bulk RNA sequencing on lung tissues from vehicle- and anti-mIL-6 Ab-treated SuHx mice (*n* = 3 *per* group). Principal Component Analysis (PCA) and sample correlation matrix revealed distinct clustering between groups, indicating a clear separation in transcriptomic profiles following IL-6 blockade (**Figure 5A-B**). Heatmap and differential gene expression (DEG) analyses further highlighted significant transcriptional differences: 101 genes were upregulated, and 23 genes were downregulated in anti-mIL-6 Ab-treated mice compared with vehicle-treated controls (**Figure 5B-C**). Pathway enrichment analysis using Gene Ontology (biological processes) and KEGG revealed that IL-6 neutralization significantly modulated immune response, inflammatory pathways, and the IL6-JAK-STAT3 signaling axis (**Figure 5D-F**). Transcription factor (TF) enrichment analysis (ChEA) identified STAT3, STAT4, and FOXP3 as critical regulators whose downstream targets were differentially expressed after IL-6 blockade (**Figure 5G-H**). FOXP3 is a master transcriptional regulator of immune tolerance, essential for regulatory T cell function and suppression of inflammation^43–45^. Notably, FOXP3 expression was suppressed in vehicle-treated SuHx mice and restored upon IL-6 neutralization, suggesting that IL-6-driven FOXP3 repression contributes to pulmonary vascular remodeling and immune dysregulation in PAH **(Figure 5G-H)**. Motif analysis of the FOXP3 locus using JASPAR identified predicted STAT4 binding sites within the FOXP3 gene, raising the possibility that STAT4, potentially in cooperation with STAT3, contributes to transcriptional repression of FOXP3 (**Figure 5I**). These findings support a model in which IL-6 promotes vascular inflammation and remodeling in PAH through STAT3/STAT4-dependent repression of FOXP3.

**Figure 5.**
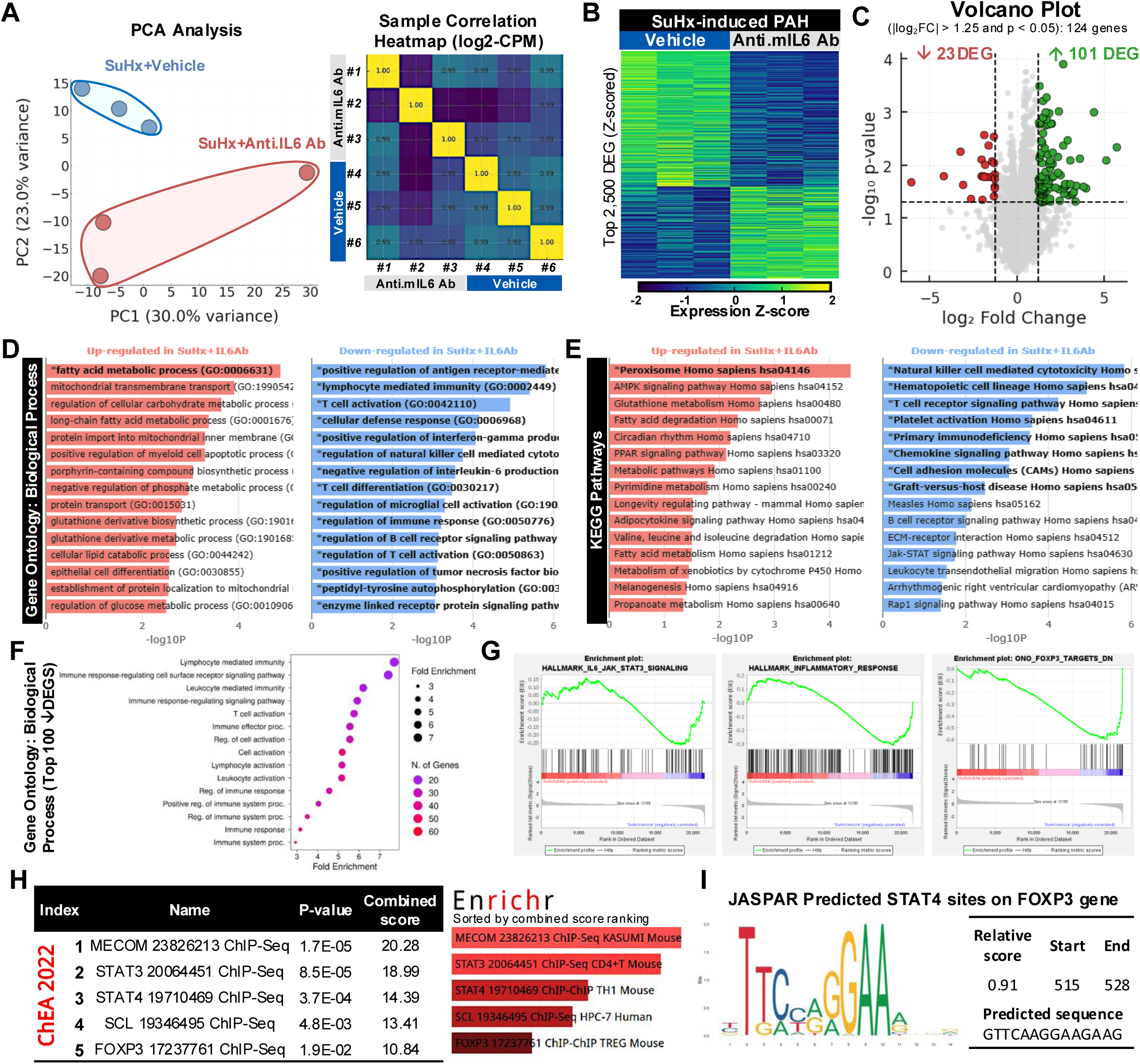
Transcriptomic and pathway analysis of anti–mouse IL-6 antibody treatment in whole lungs from the SuHx-induced PAH model. **(A-C)** Principal component analysis (PCA), heatmap, and volcano plot illustrating genes that are significantly upregulated or downregulated in SuHx mouse lungs treated with anti–mouse IL-6 neutralizing antibody (anti-mIL6 Ab) compared with vehicle-treated controls (n = 3 per group). **(D-E)** Gene Ontology and Kyoto Encyclopedia of Genes and Genomes (KEGG) pathway enrichment analyses of significantly differentially expressed genes following anti-mIL-6 Ab treatment. **(F)** Gene Set Enrichment Analysis (GSEA) demonstrating significant regulation of immune response–related gene sets in response to anti-mIL6 Ab treatment. **(G)** GSEA showing that anti-mIL6 Ab treatment in SuHx mice significantly modulates IL-6/STAT3 signaling, inflammatory response pathways, and FOXP3-mediated transcriptional programs. **(H)** Transcription factor (TF) enrichment analysis of RNA-sequencing data using ENCODE and ChEA consensus TFs from the ChIP-X database identifies STAT3, STAT4, and FOXP3 among the top 10 significantly regulated TFs. **(I)** JASPAR-based prediction of STAT4 binding sites within the FOXP3 gene locus. Corresponding P values and combined scores are shown for each predicted transcription factor binding site. Data are presented as mean ± SEM. ***P < 0.001.

### Elevated IL-6 levels are inversely associated with FOXP3 in PAH patients

To examine downstream mechanisms of IL-6 neutralization in PH, we focused on FOXP3 and STAT4. In lung homogenates from SuHx mice and MCT rats, treatment with species-specific IL-6 neutralizing antibodies increased FOXP3 expression at both the mRNA and protein levels (**Figure 6A-D**). Circulating FOXP3 levels were also increased following IL-6 blockade in both models compared with vehicle-treated groups (**Figure 6E-F**).

**Figure 6.**
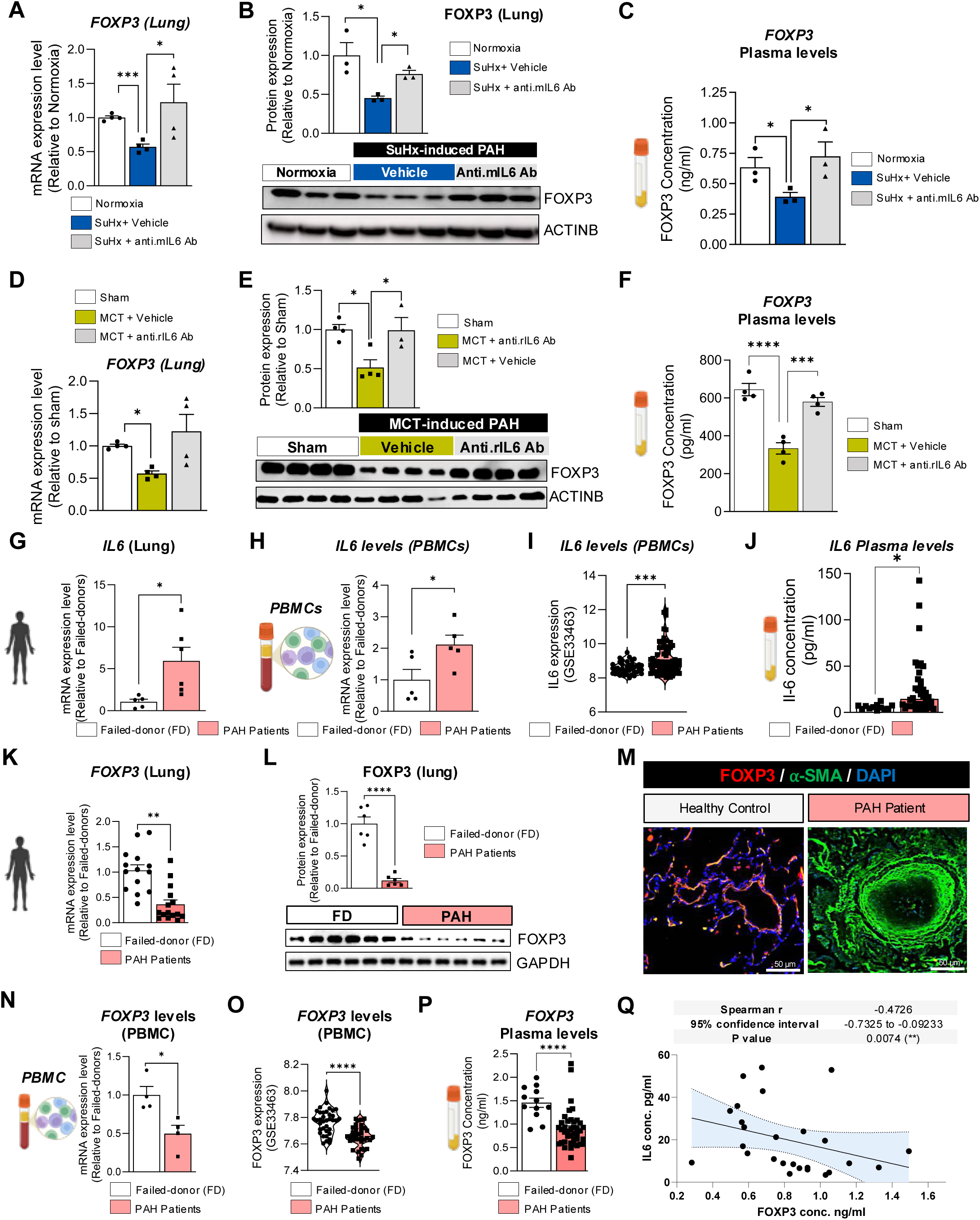
Elevated IL-6 levels are inversely associated with FOXP3 expression in experimental PAH models and human PAH. **(A-E)** FOXP3 mRNA and protein expression levels in lung tissues from control animals and MCT- or SuHx-induced PAH models treated with vehicle or species-specific IL-6 neutralizing antibodies, as assessed by RT-qPCR, immunoblotting, and immunostaining. **(F)** Plasma FOXP3 concentrations measured by ELISA in the indicated animal groups (n = 4). **(G)** IL-6 mRNA expression levels measured by RT-qPCR in lung tissues from failed donors (FD), non-PAH controls, and PAH patients (n = 5). **(H)** IL-6 expression levels in peripheral blood mononuclear cells (PBMCs) isolated from FD controls and PAH patients, reflecting chronic inflammatory activation (n = 6). **(I)** PBMC IL-6 expression levels derived from a publicly available human dataset (GSE334663). **(J)** Plasma IL-6 concentrations measured by ELISA in FD controls, non-PAH controls (n=12), and PAH patients (n = 40). **(K)** FOXP3 mRNA expression levels in lung tissues from failed donor controls assessed by RT-qPCR (n = 14-15). **(L)** Representative immunoblots (lower) and corresponding densitometric quantification (upper) of FOXP3 protein expression in whole-lung homogenates from PAH patients and FD donors (n = 6). **(M)** Representative co-immunofluorescence staining for FOXP3 (green) and endothelial cell marker CD31 (red) in lung sections from failed donors and PAH patients. Nuclei were counterstained with DAPI (blue). Scale bar = 50 μm. **(N)** IL-6 mRNA expression levels in cultured PBMCs from FD controls and PAH patients (n = 4). **(O)** PBMC IL-6 expression levels in the same human dataset (GSE334663). **(P)** Plasma FOXP3 concentrations measured by ELISA in failed donor controls (n = 12) and PAH patients (n = 40). **(Q)** Correlation analysis between IL-6 and FOXP3 expression levels across experimental models and human PAH samples. Data are presented as mean ± SEM. *P < 0.05, **P < 0.01, ***P < 0.001.

Clinically, IL-6 is elevated in the lung tissue and blood of PAH patients and correlates negatively with clinical outcomes^8,12,13,30,46–48^. Analysis of human lung samples confirmed significantly higher IL-6 mRNA levels in PAH patients compared with healthy controls (**Figure 6G & Supplementary Table 1**). These findings were corroborated in cultured PBMCs from PAH patients and in published transcriptomic datasets derived from PBMCs, which consistently showed elevated IL-6 levels (**Figure 6H-I**). Plasma IL-6 levels were also increased in PAH patients (**Figure 6J**). FOXP3 mRNA and protein levels were significantly reduced in PAH lung homogenates compared with controls (**Figure 6K-L**), as assessed by QPCR and immunoblot. Importantly, immunostaining further confirmed marked reductions in distal pulmonary vessels, particularly in the endothelial layer (**Figure 6M**). Similarly, FOXP3 mRNA was decreased in PBMCs from PAH patients and in published transcriptomic datasets (**Figure 6N-O**). Consistently, circulating FOXP3 levels were lower in patients with idiopathic PAH (IPAH) and systemic sclerosis-associated PAH (SSc-PAH) (**Supplementary Table 2)**, and were associated with increased IL-6 levels and adverse clinical outcomes (**Figure 6P-Q**). Collectively, these findings indicate that IL-6-driven FOXP3 suppression is conserved across lung tissue, PBMCs, and serum, supporting FOXP3 as a mechanistic effector and a potential predictive biomarker for response to anti-IL-6 therapy. Measurement of IL-6 and FOXP3 levels may inform patient stratification and guide precision therapeutic strategies in PAH.

### IL-6 regulates FOXP3 expression in PAH-derived PAECs via STAT3/STAT4 interaction

IL-6 is abundantly expressed not only by immune cells but also by pulmonary vascular cells, including PAECs and PASMCs ^49^. In PAH, elevated IL-6 expression in pulmonary vascular cells promotes vascular proliferation and inflammation through autocrine and paracrine signaling ^27,50^. IL-6 is also known to negatively regulate BMPR2 expression and downstream signaling via STAT3 hyperactivation^42^. Although STAT4 has minimal effects on the development of natural regulatory T cells (Tregs), it can influence FOXP3 regulation ^51^. To dissect IL-6-mediated FOXP3 regulation in pulmonary vascular cells, we first quantified IL-6 mRNA in primary PAECs and PASMCs derived from failed donor and PAH patients. IL-6 expression was increased in both PAH-PAECs and PAH-PASMCs compared with FD controls, with a greater induction in PAH-PAECs (**Figure 7A**). FOXP3 mRNA levels were reduced in both PAH-PAECs and PAH-PASMCs relative to FD cells (**Figure 7B**). In contrast, STAT4 transcripts were increased in PAH-PAECs but unchanged in PAH-PASMCs (**Figure 7C**). Similarly, immunoblot analysis validated reduced FOXP3 expression in PAH-PAECs, accompanied by increased STAT4 and pSTAT3 levels compared to FD controls **(Figure 7D & Supplementary Figure 5A-C).**

**Figure 7.**
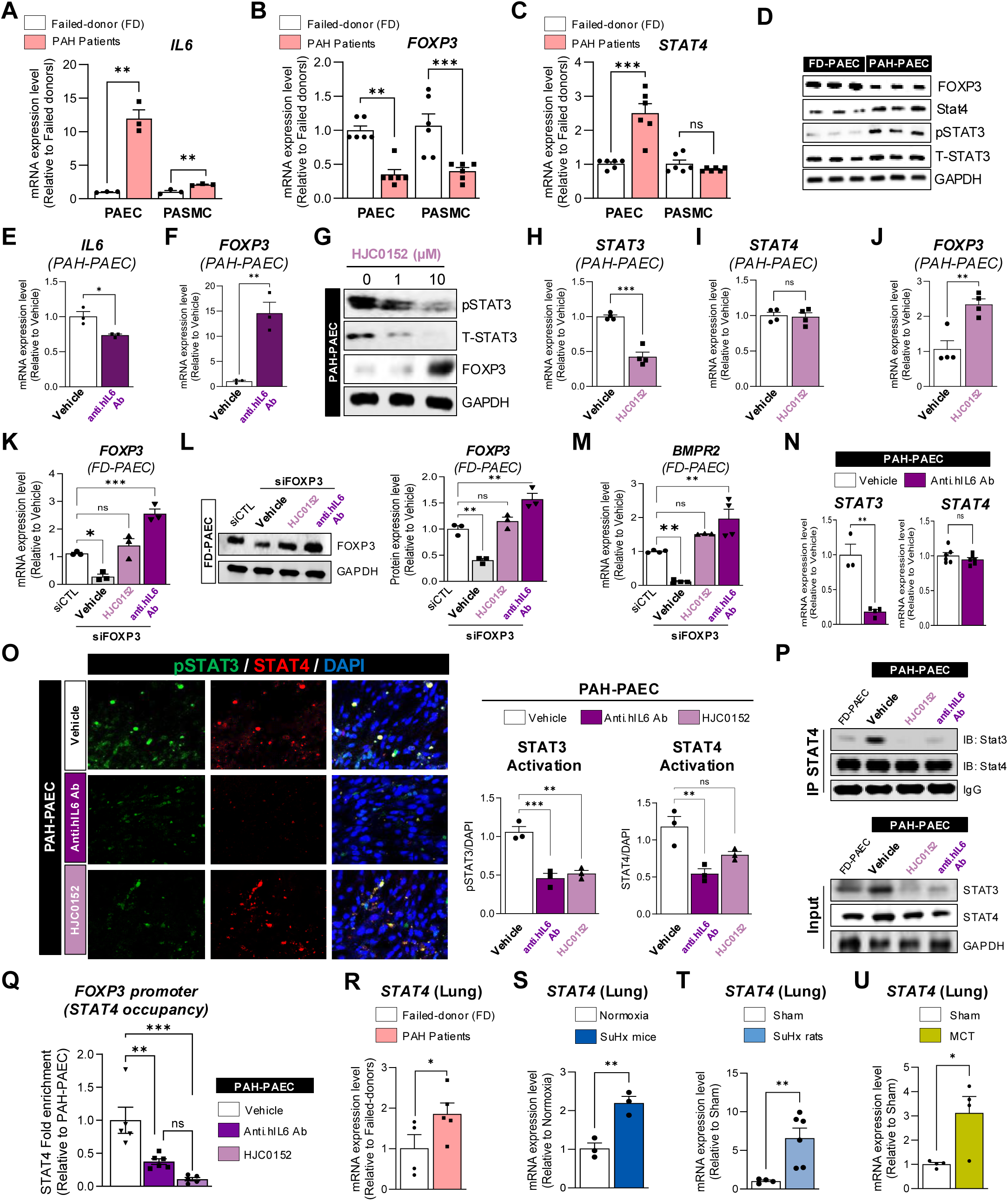
IL-6 represses FOXP3 expression in PAECs through STAT3-STAT4 interaction. **(A-C)** Relative mRNA expression levels of IL-6, FOXP3, and STAT4 measured by RT-qPCR in PAECs and PASMCs derived from failed donors (FD) and PAH patients (n = 3-6). **(D)** Representative immunoblot analysis of FOXP3, STAT4, phosphorylated STAT3 (p-STAT3), and total STAT3 in FD-PAECs and PAH-PAECs (n = 3). **(E-F)** IL-6 and FOXP3 mRNA expression levels in PAH-PAECs following 48-hour treatment with vehicle or human monoclonal anti–IL-6 antibody (anti-hIL6 Ab; 20 μM) (n = 3). **(G)** Representative immunoblot analysis of p-STAT3, total STAT3, and FOXP3 in PAH-PAECs treated for 48 hours with vehicle or increasing concentrations of the STAT3 inhibitor HJC0152. (H-J) STAT3; STAT4 and FOXP3 mRNA expression levels in PAH-PAECs following 48-hour treatment with vehicle or STAT3 inhibitor HJC0152 **(K-L)** FOXP3 and BMPR2 mRNA expression levels measured by RT-qPCR in FOXP3 siRNA–depleted FD-PAECs after 72 hours, followed by treatment with HJC0152 or anti-hIL6 Ab (20 μM) for an additional 48 hours (n = 3). **(M)** Representative immunoblot analysis of FOXP3 in FOXP3 siRNA–depleted FD-PAECs treated with HJC0152 or anti-hIL6 Ab (20 μM). Densitometric quantification normalized to GAPDH is shown (upper panel) (n = 3). **(N)** STAT3 and STAT4 mRNA expression levels assessed by RT-qPCR in PAH-PAECs following 48-hour treatment with anti-hIL6 Ab (n = 3). **(O)** Representative confocal images showing co-immunofluorescence staining of p-STAT3 (green) and STAT4 (red) in PAH-PAECs treated with HJC0152 or anti-hIL6 Ab (20 μM) for 48 hours (left). Quantification of p-STAT3 and STAT4 signals normalized to DAPI-stained nuclei is shown (right). Scale bar = 100 μm. **(P)** Immunoprecipitation of STAT4 followed by immunoblot analysis of STAT3 and STAT4 in PAH-PAECs treated with HJC0152 or anti-hIL6 Ab under the indicated conditions. **(Q)** Chromatin immunoprecipitation–qPCR (ChIP-qPCR) analysis assessing STAT4 binding to the FOXP3 promoter in PAH-PAECs treated with vehicle, HJC0152, or anti-hIL6 Ab. Promoter occupancy is expressed as fold change relative to vehicle-treated control cells. **(R-U)** STAT4 mRNA expression levels assessed by RT-qPCR in lung homogenates from failed donor controls (n = 5), idiopathic PAH (iPAH) patients (n = 5), and the indicated experimental PAH animal models (n = 3-6). Data are presented as mean ± SEM. *P < 0.05, **P < 0.01, ***P < 0.001.

To test whether IL-6 directly regulates FOXP3 in PAH-PAECs, cells were treated with a human IL-6 neutralizing antibody (anti-hIL6 Ab). IL-6 neutralization significantly reduced IL-6 expression and restored FOXP3 mRNA and protein levels (**Figure 7E-F**). Conversely, FD-PAECs stimulated with a human IL-6 agonist exhibited increased IL-6 expression and reduced FOXP3 levels, both of which were reversed by anti-hIL6 Ab treatment (**Supplementary Figure 5D-E**). Moreover, anti-hIL6 Ab treatment also reduced proliferation and migration of FD-PAECs stimulated with a human IL-6 agonist and cultured under high-serum concentrations **(Supplementary Figure 6A-B**). Mechanistically, STAT3 inhibition with HJC0152 in PAH-PAECs reduced STAT3 activation and partially restored FOXP3 protein levels (**Figure 7G & Supplementary Figure 6A-B)**. At 10 μM, HJC0152 reduced STAT3 and FOXP3 mRNA levels without affecting STAT4 expression. **(Figure 7H-J)**. In FOXP3-depleted FD-PAECs, IL-6 neutralization rescued both FOXP3 and BMPR2 expressions, whereas STAT3 inhibition produced only a partial effect, indicating that IL-6 suppresses FOXP3 primarily via STAT3 but may involve additional mechanisms (**Figure 7K-M**). We next assessed the role of STAT4 in FOXP3 regulation. Immunostaining revealed increased STAT3 and STAT4 and reduced FOXP3 expression in PAH-PAECs (**Supplementary Figure 7A**). IL-6 blockade reduced STAT3 expression without affecting STAT4, as confirmed by immunofluorescence (**Figure 7N-O**). Co-immunoprecipitation revealed increased STAT3–STAT4 interaction in PAH-PAECs, which was attenuated by the STAT3 inhibitor HJC0152 and by anti-hIL-6 neutralization (**Figure 7P**). Given that in silico analysis of the FOXP3 promoter identified predicted STAT4 binding motifs **(Figure 5I)**, we examined STAT4 occupancy by ChIP–qPCR. STAT4 enrichment at the FOXP3 promoter was reduced following STAT3 inhibition or IL-6 neutralization in PAH-PAECs **(Figure 7Q)**. Together, these findings support a cooperative mechanism in which STAT3–STAT4 interaction contributes to repression of FOXP3 expression in PAH-PAECs. Finally, STAT4 mRNA levels were significantly increased in lung tissue from patients with PAH **(Figure 7R)** and in lungs from experimental PH rodent models **(Figure 7S-U)**, supporting a pathological role for STAT4 signaling in PH. Together, these findings further support a model in which IL-6 suppresses FOXP3 expression in PAH-derived PAECs predominantly via STAT3 activation, with modulation by STAT3/STAT4 interaction, thereby contributing to endothelial dysfunction and pulmonary vascular remodeling.

### IL-6 regulates FOXP3 through DNA methylation in PAH

In addition to STAT-dependent transcriptional repression, we investigated whether IL-6 modulates FOXP3 expression through epigenetic mechanisms. In silico analysis of the *FOXP3 p*romoter identified multiple CpG sites and islands, suggesting also a susceptibility to methylation-dependent repression (**Figure 8A**). In FD-PAECs, IL-6 stimulation increased DNMT1 expression, whereas 5-aza-2′-deoxycytidine (5-Aza), a DNA methyltransferase inhibitor used to reduce DNMT1-mediated DNA methylation, reduced DNMT1 and IL-6 levels (**Figure 8B-C**). Interestingly, we found that STAT3 inhibition similarly reduced DNMT1 mRNA expression (**Figure 8D**), consistent with previous studies demonstrating that direct regulation of DNMT1 by STAT3^52,53^. This downregulation of DNMT1 may lead to the demethylation of the *FOXP3* promoter, thereby restoring its expression. Indeed, the combined treatment with anti-hIL6 Ab and 5-Aza markedly decreased *FOXP3* promoter methylation in the presence of IL6 agonist (**Figure 8E**) and significantly increased FOXP3 mRNA and protein expression in FD-PAECs (**Figure 8F**).

**Figure 8.**
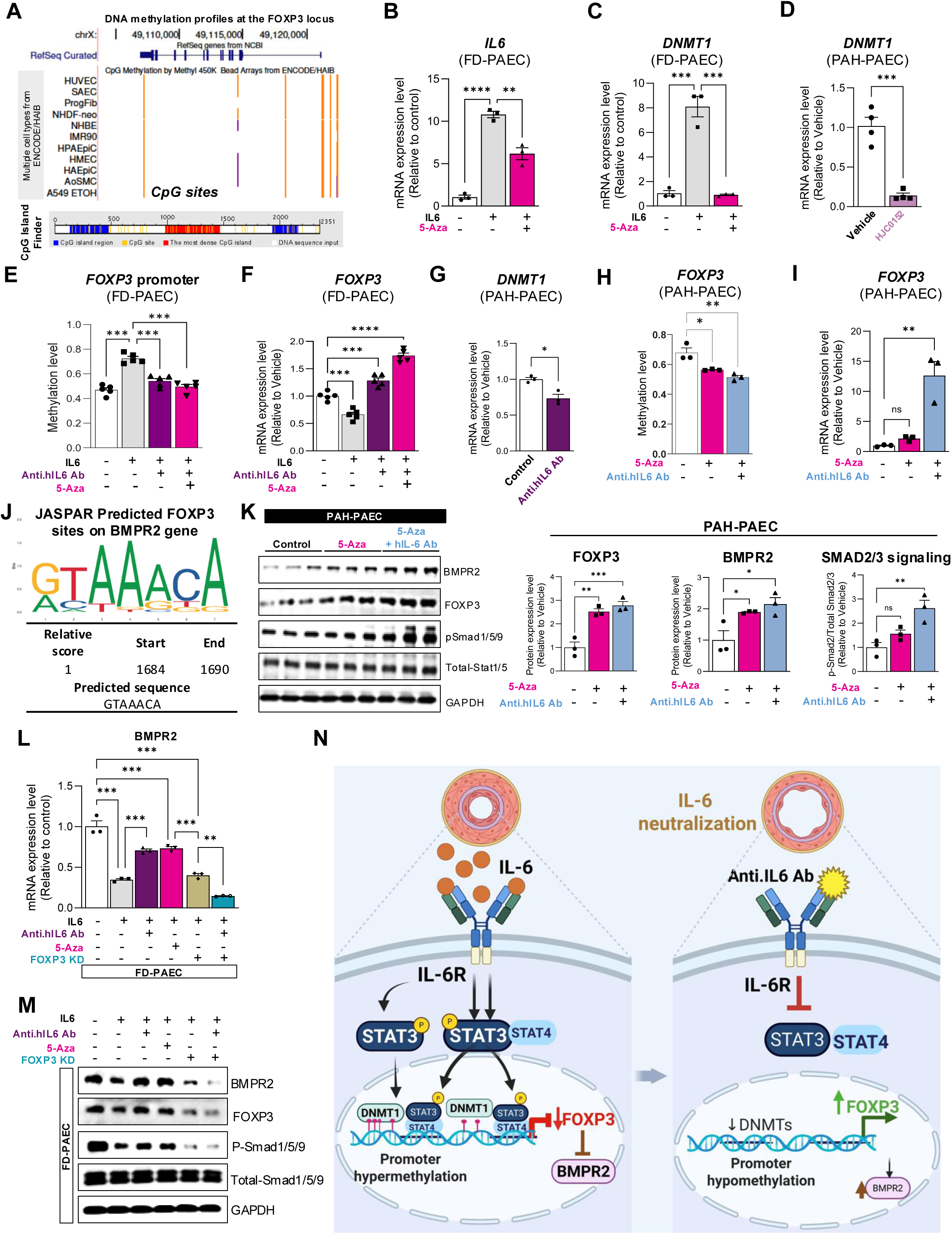
IL-6 suppresses FOXP3 expression through DNMT1-mediated promoter methylation. **(A)** UCSC Genome Browser view of the FOXP3 promoter region showing ENCODE DNA methylation data (upper panel) and CpG island prediction using the CpG Island Finder tool (lower panel). CpG islands and methylation sites are indicated in blue and red, respectively. **(B-C)** IL-6 and DNMT1 mRNA expression levels assessed by RT-qPCR in failed donor-derived PAECs (FD-PAECs) treated with recombinant IL-6 in the presence or absence of the DNA methyltransferase inhibitor 5-azacytidine (5-Aza; 5 μM) for 48 hours (n = 3). **(D)** DNMT1 mRNA expression levels assessed by RT-qPCR in PAH-PAECs in the presence or absence of the STAT3 inhibitor HJC0152 for 48 hours (n = 3). **(E)** Methylation-specific qPCR (MSP) analysis identifying methylated CpG sites within the *FOXP3* promoter region in FD-PAECs treated with IL-6 ± 5-Aza ± anti-rabbit IL-6 neutralizing antibody (anti-hIL6 Ab) for 48 hours (n = 4). **(F)** FOXP3 mRNA expression levels measured by RT-qPCR under the same treatment conditions as in (E) (n = 4). **(G)** DNMT1 mRNA expression levels assessed by RT-qPCR in PAH-PAECs treated with vehicle or anti-hIL6 Ab for 48 hours (n = 3). **(H)** MSP qPCR analysis of *FOXP3* promoter methylation in PAH-PAECs treated with 5-Aza ± anti-hIL6 Ab (n = 5). **(I)** FOXP3 mRNA expression levels in PAH-PAECs treated with vehicle or 5-Aza ± anti-hIL6 Ab for 48 hours (n = 4). **(J)** JASPAR-based prediction of conserved FOXP3 binding sites within the BMPR2 gene locus. **(K)** Representative immunoblot analysis of BMPR2, FOXP3, phosphorylated SMAD1/5/9 (p-SMAD1/5/9), and total SMAD1/5/9 in PAH-PAECs treated with 5-Aza (5 μM) in the presence or absence of anti-hIL6 Ab (20 μM). Densitometric quantification is shown (right) (n = 3). **(L)** BMPR2 mRNA expression levels measured by RT-qPCR in FOXP3 siRNA-depleted FD-PAECs treated with IL-6 ± 5-Aza ± anti-hIL6 Ab for 48 hours (n = 3). **(M)** Representative immunoblots of BMPR2, FOXP3, p-SMAD1/5/9, total SMAD1/5/9, and GAPDH (loading control) in FOXP3 siRNA-depleted FD-PAECs treated with IL-6 ± 5-Aza ± anti-hIL6 Ab for 48 hours (n = 3). **(N)** Schematic proposed a working model summarizing the dual mechanisms by which IL-6 represses FOXP3 via STAT3/STAT4 interaction and DNMT1-mediated promoter methylation, thereby suppressing BMPR2, promoting vascular inflammation, and driving pulmonary arterial remodeling. Data are presented as mean ± SEM. *P < 0.05, **P < 0.01, ***P < 0.001.

In PAH-PAECs, 5-Aza alone reduced DNMT1 expression, and co-treatment with anti-hIL6 Ab synergistically reduced *FOXP3* promoter methylation and robustly restored FOXP3 mRNA expression levels (**Figure 8H-I**). To assess the functional relevance of FOXP3, we analyzed the transcription factor binding site (TFBS) of the *BMPR2* promoter, which revealed a predicted FOXP3 binding motif (**Figure 8J**). Interestingly, co-treatment with anti-hIL6 Ab and 5-Aza significantly restored FOXP3 and BMPR2 protein levels, as well as SMAD1/5/9 downstream signaling in PAH-PAECs (**Figure 8K**). Consistently, FOXP3 knockdown in FD-PAECs reduced BMPR2 expression following IL-6 stimulation, whereas 5-Aza, alone or combined with anti–hIL-6 Ab, restored FOXP3, BMPR2, and downstream SMAD1/5/9 signaling (**Figure 8L-M**).

Collectively, these findings support a model in which IL-6 suppresses FOXP3 through both STAT3/STAT4-dependent mechanisms and DNMT1-mediated promoter methylation. Restoration of FOXP3 reduces inflammatory signaling and rescues BMPR2 expression, supporting a role for IL-6 blockade in limiting pulmonary vascular remodeling in PAH (**Figure 8N**).

## DISCUSSION

The most compelling novelty of our study lies in the identification of FOXP3 as a pivotal downstream effector of IL-6 signaling in PAH. For the first time, we establish that IL-6-mediated suppression of FOXP3 in PAECs and PBMCs serves as the definitive mechanistic link between chronic inflammatory signaling and pathological vascular remodeling. While FOXP3 is traditionally recognized as the master regulator of immune tolerance in T cells, our data reveals its critical, previously unrecognized role in maintaining endothelial homeostasis. Mechanistically, we demonstrate that IL-6 drives this suppression through a sophisticated dual-regulatory program: transcriptional repression via a non-canonical STAT3/STAT4 interaction and epigenetic silencing through DNMT1-mediated promoter methylation. The discovery that STAT4, typically essential for Treg development, is co-opted to repress FOXP3 in the PAH lung represents a significant shift in our understanding of cytokine-driven vascular injury. By showing that IL-6 neutralization restores FOXP3 expression and reverses endothelial dysfunction, we define the IL-6/STAT3-STAT4/FOXP3 axis as a novel therapeutic paradigm for restoring vascular integrity in PAH.

FOXP3 has been shown as a master regulator of Tregs and is essential for maintaining immune tolerance and suppressing pro-inflammatory pathways^44,45^. Previous studies have implicated FOXP3 dysfunction in immune dysregulation in idiopathic and connective tissue-associated PAH ^54^. Our data indicate that IL-6 neutralization restores FOXP3 expression in PAH-PAECs *in vitro* and in treated animals, highlighting a previously unrecognized endothelial target of IL-6/FOXP3 and a potential mediator of vascular homeostasis.

Mechanistically, we revealed that IL-6 orchestrates this suppression through a sophisticated dual-regulatory program. First, we identify a non-canonical STAT3/STAT4 interaction and epigenetic silencing via DNMT1-mediated methylation of the FOXP3 promoter. Although STAT4 is traditionally associated with Th1 differentiation and considered indispensable for Treg development, we show that STAT4 expression and promoter occupancy are increased in PAH lungs and PAECs. ChIP-qPCR analysis showed reduced STAT4 occupancy at the FOXP3 promoter following IL-6 neutralization or STAT3 inhibition, supporting a role for STAT3-STAT4 interaction in the regulation of FOXP3 in PAECs. This non-canonical mechanism represents a novel paradigm by which inflammatory cytokines modulate endothelial transcriptional programs classically associated with immune tolerance.

Previous studies have shown that IL6 activates the STAT3 signaling pathway, which then upregulates the miR-17/92 cluster, which downregulates BMPR2 expression ^42^. In addition to STAT3/STAT4, our findings identify epigenetic regulation as a complementary mechanism by which IL-6 suppresses FOXP3. IL-6 increases DNMT1 expression in PAH-PAECs, leading to hypermethylation of the *FOXP3* promoter and sustained transcriptional repression. Our findings establish a direct link between proinflammatory STAT3 signaling and the epigenetic silencing of FOXP3, mediated by DNMT1. STAT3 is well-known for its role in Th17 differentiation; its ability to actively suppress the Treg program via DNA methylation provides a dual mechanism for shifting the T-cell balance^52,55,56^. Consistent with previous studies conducted in malignant T cells^52^, our data show that STAT3 acts as a transcriptional driver of DNMT1. By maintaining elevated DNMT1 expression and promoting interaction with STAT4, p-STAT3 contributes to sustained hypermethylation of the FOXP3 promoter, leading to stable transcriptional repression. This mechanism is particularly relevant in PAH, where persistent STAT3 activation may impair PAEC function by limiting FOXP3 accessibility and expression. Previously, studies have shown that the recruitment of DNMT1 by STAT3 to the *FOXP3* locus appears to be a key event in maintaining immune imbalance in conditions such as latent autoimmune diabetes in adults and other autoimmune pathologies ^53^. This is further complicated by the fact that DNMT1 is indispensable for maintaining the functional identity of existing Tregs; its loss leads to lethal autoimmunity due to the “melting” of Treg-specific epigenetic marks. In our study, this axis suggests that DNMT1 is not merely a global maintenance enzyme in this context, but a specific tool recruited by IL6/STAT3 to modulate lung EC plasticity. In these contexts, hyperactivated STAT3 binds to STAT4 and ensures that the *FOXP3* promoter remains hypermethylated through DNMT1, preventing the binding of essential transcription factors and locking the PAECs in a dysfunctional state. Furthermore, pharmacological inhibition of DNMT1 or IL-6 neutralization restores FOXP3 expression and rescues BMPR2 signaling, establishing FOXP3 as a functional upstream regulator of BMPR2 in PAECs. This mechanistic link connects inflammation, epigenetic remodeling, and loss of BMPR2 signaling-all central features of PAH pathobiology.

These molecular insights are supported by robust in vivo evidence. Unlike previous studies that focused primarily on pulmonary hemodynamics, our study integrates structural, functional, molecular, and cellular assessments. Using dual-model validation in SuHx mice and MCT rats, we show that IL-6 blockade markedly ameliorates established pathology, reducing RV pressures and reversing maladaptive remodeling. Cardiac MRI and hemodynamic analyses consistently demonstrate that neutralizing IL-6 restores RV function (RVEF, TAPSE) and reduces interstitial fibrosis. These findings underscore IL-6 as the central molecular switch linking pulmonary vascular injury to RV failure. Collectively, these findings underscore IL-6 as a central molecular driver linking vascular and RV remodeling and highlight its neutralization as a potential switch in disease progression.

Furthermore, our data provides a critical mechanistic lens through which to view the discrepancies between preclinical success and recent clinical trials. The TRANSFORM-UK trial (NCT02676947) evaluating Tocilizumab in PAH patients failed to demonstrate significant improvements in pulmonary vascular resistance or exercise capacity ^31–33,57^. While our study utilized direct ligand neutralization in early-stage disease to achieve robust reversal of pathology, the clinical divergence may also stem from the timing of intervention and the specific cellular targets involved. Our discovery of the FOXP3/BMPR2 link suggests that therapeutic efficacy may depend on the ability to overcome the “epigenetic mask” established by chronic IL-6 exposure. In summary, we demonstrate that IL-6/STAT3 signaling recruits DNMT1 and STAT4 to selectively silence FOXP3, thereby stripping the pulmonary endothelium of its protective phenotypic identity. By identifying FOXP3 as a guardian of endothelial BMPR2 signaling, our study provides a compelling rationale for targeting this axis to halt, and potentially reverse, the progression of PAH.

### Clinical strategies to measure IL-6 and FOXP3 as a biomarker in PAH

Importantly, our work also carries translational implications for biomarker development. While IL-6 reflects systemic inflammatory burden, FOXP3 integrates downstream transcriptional and epigenetic consequences of IL-6 signaling in PAECs, including the restoration of BMPR2. Our findings suggest that measuring FOXP3, alone or in combination with IL-6, could provide a mechanistically informed biomarker of vascular target engagement. Assessment of *FOXP3* promoter methylation in PBMCs offers a minimally invasive, scalable approach to monitor the effectiveness of IL-6-targeted therapy. These findings raise the possibility that elevated IL-6 alone may not be sufficient to predict therapeutic response. Patients with high IL-6 levels but preserved FOXP3 expression may represent a distinct subgroup less dependent on this pathway and therefore less likely to benefit from IL-6-targeted therapies. Integrating IL-6 and FOXP3 profiling could support a precision medicine approach by guiding patient stratification and informing therapeutic selection.

### Study limitations

Despite strong preclinical findings, several limitations should be considered. Human samples were derived from cross-sectional cohorts, often representing advanced disease, which limit conclusions about causality. In addition, PBMC analyses reflect a heterogeneous cell population, which may obscure cell-specific contributions to FOXP3 regulation. Sample sizes in human cohorts remain modest and may not capture variability across PAH subtypes. Finally, clinical translation remains uncertain, as the only completed trial of IL-6 blockade in PAH (TRANSFORM-UK) was negative, although ongoing studies such as the SATISFY trial may provide additional insight^31,58^.

## CONCLUSION

Collectively, our study positions FOXP3 as a central molecular hub integrating inflammatory, transcriptional, and epigenetic signals in the pulmonary vasculature, where its suppression via IL-6-driven STAT3/STAT4 interaction and DNMT1-mediated promoter hypermethylation drives endothelial dysfunction and vascular remodeling. This dual mechanism silencing of FOXP3 subsequently triggers the downregulation of BMPR2 signaling, a hallmark of PAH pathogenesis, thereby establishing a direct link between pro-inflammatory cytokine elevation and the loss of pulmonary vascular homeostasis. While translational applicability is currently constrained by the cross-sectional nature of end-stage human lung samples and the cellular heterogeneity inherent in systemic PBMCs, our results demonstrate that restoring FOXP3, whether through IL-6 blockade, STAT3 inhibition, or epigenetic modulation with DNMT1-targeting therapies such as 5-Aza, can effectively rescue BMPR2 expression and attenuate disease progression. These findings provide a robust mechanistic rationale for IL-6-targeted therapies and distinguish FOXP3 as both a promising, mechanistically informed biomarker for monitoring treatment efficacy and a novel focal point for personalized molecular diagnostics and therapeutic strategies in PAH.

## Authors contributions

P.K, K.J., A.G., C.G.T, T.D.J, M.M.O, MM, A.A, Y.E., and S.Z. performed the experiments and collected and analyzed the data. P.M.H. and H.P. contributed to the study of design for humans and provided the essential experimental resources. H.P, Y.S and P.M.H contributed to manuscript editing and critical revision of intellectual content. P.K., M.B., and L.H. conceived and designed the study, supervised the research, performed the experiments, analyzed and interpreted the data, and wrote the manuscript. All authors have reviewed and approved the final manuscript.

## Conflict of Interest declaration

The authors declare that they have no affiliations with or involvement in any organization or entity with any financial interest in the subject matter or materials discussed in this manuscript.

## Supporting information

Expanded Methods

## Acknowledgments

Not applicable. All the contributions to this article are limited to the authors listed in the manuscript.

## Funding

This study was supported by the following grants: National Institutes of Health (NIH)/National Heart, Lung, and Blood Institute (NHLBI) R01HL172043, R01HL158998-01A1, R01HL173203-01, NIH/National Center for Advancing Translational Sciences R03TR004673 (to L. Hadri); American Lung Association Innovation Award 1056600 and American Heart Association Award (to L. Hadri); and NIH/NHLBI K01HL159038, NIH R25HL146166, American Heart Association 24CDA1269532, TPA, and American Thoracic Society Unrestricted Grant 23-24U1 (to M. Bisserier).

## Supplementary Tables

**Supplementary Table 1.**
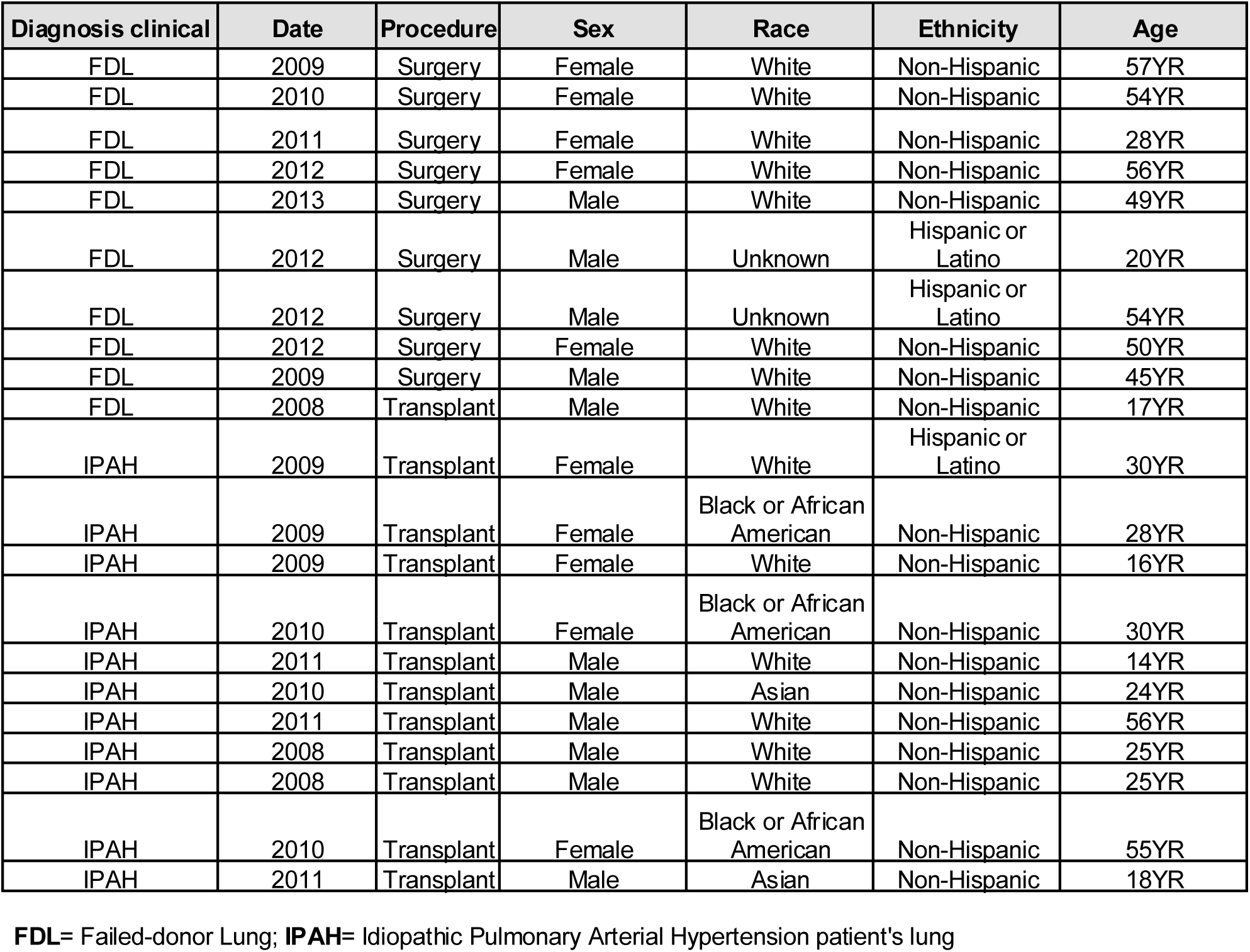
Clinical characteristics of human subjects. Demographic and clinical data of failed donor controls and PAH patients, including age, sex, diagnosis, hemodynamics, and treatment status, were available.

| Diagnosis clinical | Date | Procedure | Sex | Race | Ethnicity | Age |
| --- | --- | --- | --- | --- | --- | --- |
| FDL | 2009 | Surgery | Female | White | Non-Hispanic | 57YR |
| FDL | 2010 | Surgery | Female | White | Non-Hispanic | 54YR |
| FDL | 2011 | Surgery | Female | White | Non-Hispanic | 28YR |
| FDL | 2012 | Surgery | Female | White | Non-Hispanic | 56YR |
| FDL | 2013 | Surgery | Male | White | Non-Hispanic | 49YR |
| FDL | 2012 | Surgery | Male | Unknown | Hispanic or Latino | 20YR |
| FDL | 2012 | Surgery | Male | Unknown | Hispanic or Latino | 54YR |
| FDL | 2012 | Surgery | Female | White | Non-Hispanic | 50YR |
| FDL | 2009 | Surgery | Male | White | Non-Hispanic | 45YR |
| FDL | 2008 | Transplant | Male | White | Non-Hispanic | 17YR |
| IPAH | 2009 | Transplant | Female | White | Hispanic or Latino | 30YR |
| IPAH | 2009 | Transplant | Female | Black or African American | Non-Hispanic | 28YR |
| IPAH | 2009 | Transplant | Female | White | Non-Hispanic | 16YR |
| IPAH | 2010 | Transplant | Female | Black or African American | Non-Hispanic | 30YR |
| IPAH | 2011 | Transplant | Male | White | Non-Hispanic | 14YR |
| IPAH | 2010 | Transplant | Male | Asian | Non-Hispanic | 24YR |
| IPAH | 2011 | Transplant | Male | White | Non-Hispanic | 56YR |
| IPAH | 2008 | Transplant | Male | White | Non-Hispanic | 25YR |
| IPAH | 2008 | Transplant | Male | White | Non-Hispanic | 25YR |
| IPAH | 2010 | Transplant | Female | Black or African American | Non-Hispanic | 55YR |
| IPAH | 2011 | Transplant | Male | Asian | Non-Hispanic | 18YR |

**FDL**= Failed-donor Lung; **IPAH**= Idiopathic Pulmonary Arterial Hypertension patient's lung

**Supplementary Table 2.** Clinical characteristics of human serum samples, including demographic, structural, clinical, functional, echocardiographic, hemodynamic characteristics, and immune markers.

**FDL**= Failed-donor Lung; **IPAH**= Idiopathic Pulmonary Arterial Hypertension patient's lung
| Variable | IPAH patients<br>(n=7) 33.34% | SSc-PAH patients<br>(n=14) 66.66% | P value |
| --- | --- | --- | --- |
| **Demographic Characteristics** |  |  |  |
| Age at PH diagnosis, yr | 55.43 ± 12.2 | 59.43 ± 12.6 | 0.496 |
| Female | 5 (71.4) | 12 (85.7) | 0.574 |
| Race |  |  | 0.687 |
| White | 6 (85.7) | 12 (85.7) |  |
| African American | 1 (14.3) | 1 (7.1) |  |
| Other | 0 (0.0) | 1 (7.1) |  |
| Deaths | 0 (0.0) | 9 (64.3) | 0.007 |
| **PAH Therapy** |  |  |  |
| Received PAH therapy | 5 (71.4) | 14 (100) | 0.100 |
| Number of PAH Therapies* |  |  | 0.397 |
| ≤1 | 4 (57.1) | 3 (42.9) |  |
| ≥2 | 5 (35.7) | 9 (64.3) |  |
| **Structural, Clinical, Functional, Echocardiographic, and Hemodynamic Characteristics at the Time of Diagnosis** |  |  |  |
| **Structural** |  |  |  |
| RV Mass, g | 29 (22-39) | 27.5 (20.5-34.3) | 0.397 |
| LV Mass, g | 100.57 ± 20.6 | 90.21 ± 22.7 | 0.313 |
| RVESV, mL | 80 (68-155) | 85 (54.0-114.3) | 1.000 |
| RVEDV, mL | 177 (141-246) | 157 (125-185) | 0.659 |
| **Clinical and Functional** |  |  |  |
| WHO FC |  |  | 0.278 |
| FC I | 2 (28.6) | 2 (14.3) |  |
| FC II | 4 (57.1) | 5 (35.7) |  |
| FC III | 1 (14.3) | 7 (50.0) |  |
| FC IV | 0 (0.0) | 0 (0.0) |  |
| 6MWD, m | 449.43 ± 134.8 | 389.07 ± 114.2 | 0.332 |
| NT-proBNP, ng/L | 145 (64-326) | 365 (196-965) | 0.361 |
| **Echocardiographic** |  |  |  |
| PASP, mm Hg | 65 (46-101) | 50 (39.5-60) | 0.659 |
| LVEF, % | 58.86 ± 6.1 | 59.64 ± 5.4 | 0.778 |
| RVEF, % | 45.43 ± 10.8 | 48.64 ± 10.9 | 0.533 |
| VMI | 0.35 (0.23-0.36) | 0.30 (0.26-0.39) | 0.659 |
| RVSP, mmHg | 63 (44-94) | 49 (38.5-59.3) | 0.659 |
| RVDP, mm Hg | 8 (5-10) | 8 (4.3-12.3) | 1.000 |
| Right Ventricular-Pulmonary ..Arterial Coupling , E<sub>es</sub>/E<sub>a</sub> | 0.84 (0.54-2.03) | 0.90 (0.55-1.14) | 1.000 |
| **Hemodynamic** |  |  |  |
| PCWP, mm Hg | 10.14 ± 2.3 | 9.43 ± 4.2 | 0.617 |
| RAP, mm Hg | 7 (4-8) | 4.5 (3-11) | 0.397 |
| mPAP, mm Hg | 37 (30-65) | 30.5 (24.3-35.8) | 0.659 |
| PADP, mm Hg | 21 (21-44) | 17.5 (14.0-22.8) | 1.000 |
| PVR, WU | 5.12 (2.9-13.4) | 4.72 (2.77-6.17) | 0.659 |
| CO, L/min | 4.77 ± 1.2 | 4.74 ± 0.8 | 0.949 |
| **Immune Markers** |  |  |  |
| IL-6 concentration, pg/mL | 14.59 (3.92-90.87) | 12.26 (6.10-45.82) | 1.000 |
| FOXP3 concentration, ng/mL | 1.03 (0.83-1.49) | 0.82 (0.61-0.93) | 0.183 |

**Abbreviations:** 6MWD = 6-minute walking distance; CO = Cardiac output; E<sub>a</sub> = Effective arterial elastance; E<sub>es</sub> = End-systolic elastance; IPAH = Idiopathic pulmonary arterial hypertension; LV mass = Left ventricular mass; LVEF = Left ventricular ejection fraction; mPAP = Mean pulmonary artery pressure; NT-proBNP = N-terminal prohormone of brain natriuretic peptide; PADP = Pulmonary arterial diastolic pressure; PASP = Pulmonary arterial systolic pressure; PAH = Pulmonary arterial hypertension; PCWP = Pulmonary capillary wedge pressure; PH = Pulmonary hypertension; PVR = Pulmonary vascular resistance; RAP = Right atrial pressure; RVDP = Right ventricular diastolic pressure; RVEDV = Right ventricular end-diastolic volume; RVESV = Right ventricular end-systolic volume; RV mass = Right ventricular mass; RVEF = Right ventricular ejection fraction; RVSP = Right ventricular systolic pressure; SSc-PAH = Systemic sclerosis-associated pulmonary arterial hypertension; VMI = Ventricular mass index; WHO FC = World Health Organization functional class.
For categorical variables (e.g., race and deaths), data are presented as percentages. For continuous, normally distributed variables (e.g., PCWP and LV mass), data are presented as mean ± SD. For continuous, skewed variables (e.g., NT-proBNP and RAP), data are presented as median (lower quartile Q1 – upper quartile Q3).
\*Any therapy of the following: diuretic, calcium channel blocker, prostacyclin analogue, endothelin receptor antagonist, or phosphodiesterase type 5 inhibitor

**Supplementary Table 3.** Cardiac magnetic resonance imaging-derived right ventricular parameters in MCT-treated rats. Quantitative cMRI measurements of right ventricular structure and function in sham rats and MCT-treated rats receiving vehicle or anti-rat IL-6 neutralizing antibody, including RVLS, RVEDV, RVESV, RVSV, RVEF, TAPSE, and RV wallthickness. Data are presented as mean ± SEM.

|  | Sham (Mean±SD)<br>(n=4) | MCT+Vehicle (Mean±SD)<br>(n=4) | MCT+anti.rIL6 Ab<br>(Mean±SD) (n=4) |
| --- | --- | --- | --- |
| RVLS (%) | 23.1775±1.2784 | 18.01±1.553855 | 14.41±2.018667 |
| RVEDV (mL) | 0.425±0.059722 | 0.595±0.019149 | 0.5475±0.083417 |
| **RVESV (mL)** | 0.14±0.034641 | **0.375±0.058023\*** | 0.28±0.054467 |
| RVSV (mL) | 0.285±0.034157 | 0.22±0.054772 | 0.275±0.041231 |
| RVEF (%) | 67.38±3.670776 | 36.795±9.3911749 | 50.4775±5.463719 |
| **TAPSE (mm)** | 4.30075±0.289282 | **2.9765±0.3742606\*** | 2.1785±0.3682757 |
| **RV thickness** | 0.41375±0.05711 | **0.98225±0.192941** | 0.69375±0.166925 |
| **RV/LV ratio** | 0.48±0.084927 | **1.87577±0.442217** | 1.2482±0.393611 |
| **RV diameter** | 2.521±0.466683 | **5.91225±0.7301056** | 4.8175±1.3754719 |
| **LV diameter** | 5.16125±0.527705 | **3.98425±0.837905** | 3.296±0.96123 |

**RVLS**= Right Ventricular Longitudinal Strain; **RVEDV**= Right Ventricular End-Diastolic Volume; **RVESV**= Right Ventricular End-Systolic Volume; **RVSV**= Right Ventricular Stroke Volume; **RVEF**= Right Ventricular Ejection Fraction; **TAPSE**= Tricuspid Annular Plane Systolic Excursion; **RV thickness**= Right Ventricular thickness; **RV/LV ratio**= Right Ventricular/Left Ventricular Size Ratio; **RV diameter**= Right Ventricular Diameter; **LV diameter**= Left Ventricular Diameter

**Supplementary Table 4.** Left ventricular structure and function in MCT-treated rats. cMRI-derived left ventricular parameters demonstrating preserved LV dimensions and function across sham, MCT-vehicle, and anti-rIL6 Ab-treated groups, indicating RV-specific therapeutic effects.

|  | Sham<br>(Mean±SD) (n=4) | MCT+Vehicle (Mean±SD)<br>(n=4) | MCT+anti.rIL6 Ab<br>(Mean±SD) (n=4) |
| --- | --- | --- | --- |
| LV Mass | 0.56±0.042426 | 0.5225±0.122848 | 0.485±0.06455 |
| LVEDV | 0.61±0.055976 | 0.425±0.129228 | 0.4275±0.123929 |
| LVESV | 0.205±0.031091 | 0.15±0.037417 | 0.14±0.053541 |
| LVSV | 0.4±0.040825 | 0.275±0.092916 | 0.2875±0.080571 |
| LVEF | 65.7275±3.21501 | 64.2275±2.216008 | 67.6425±5.797042 |
| MAPSE | 2.832±0.565318 | 2.377±0.264629 | 2.311±0.312115 |

**LV MASS:** Left Ventricular Mass; **LVEDV=** Left Ventricular End-Diastolic Volume; **LVESV=** Left Ventricular End-Systolic Volume; **LVSV=** Left Ventricular Stroke Volume; **LVEF=** Left Ventricular Ejection Fraction; **MAPSE=** Mitral Annular Plane Systolic Excursion

**Supplementary Table 5.** cMRI-derived right ventricular parameters in SuHx mice. Longitudinal cMRI assessment of RV structure and function in normoxic controls and SuHx mice treated with vehicle or anti–mouse IL-6 neutralizing antibody, including RV mass, RVEF, TAPSE, RVSV, and RVESV.

|  | Normoxia (Mean±SD)<br>(n=4) | SuHx+Vehicle<br>(Mean±SD) (n=6) | SuHX+anti.mIL6 Ab<br>(Mean±SD) (n=6) |
| --- | --- | --- | --- |
| RVEDV (mL) | 0.0575±0.009574 | 0.0483±0.009832 | 0.056±0.010328 |
| RVESV (mL) | 0.0225±0.005 | 0.025±0.005477 | 0.02±0.006325 |
| **RVSV (mL)** | 0.035±0.005774 | **0.021±0.004082\*** | **0.035±0.008367\*\*** |
| **RVEF (%)** | 63.845±3.962815 | **48.118±3.689783\*\*\*\*** | **64.41±4.09959\*\*\*\*** |
| **TAPSE (mm)** | 2.1795±0.293685 | **1.447±0.382458\*** | **2.005±0.27171\*** |
| **RV thickness** | 0.18375±0.040991 | **0.312±0.020906\*\*\*\*** | **0.232±0.021757\*\*\*** |
| RV/LV ratio | 0.535±0.043865 | 0.643±0.086394 | 0.541±0.056599 |
| RV diameter | 2.15625±0.232024 | 2.113±0.224319 | 2.0985±0.293377 |
| **LV diameter** | 4.021±0.228868 | **3.302±0.270908\*\*** | **3.866±0.219379\*\*** |

**RVEDV**= Right Ventricular End-Diastolic Volume; **RVESV**= Right Ventricular End-Systolic Volume; **RVSV**= Right Ventricular Stroke Volume; **RVEF**= Right Ventricular Ejection Fraction; **TAPSE**= Tricuspid Annular Plane Systolic Excursion; **RV thickness**= Right Ventricular thickness; **RV/LV ratio**= Right Ventricular/Left Ventricular Size Ratio; **RV diameter**= Right Ventricular Diameter; **LV diameter**= Left Ventricular Diameter

**Supplementary Table 6.**
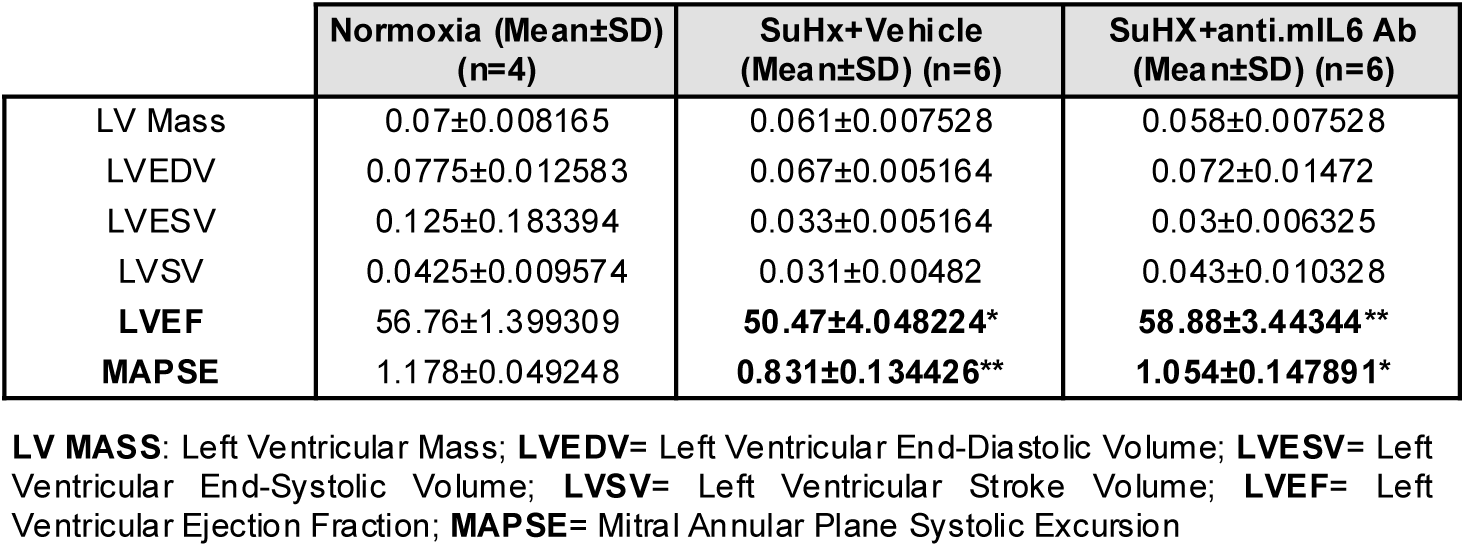
Left ventricular parameters in SuHx mice. Left ventricular structure and functional parameters assessed by cMRI in normoxic and SuHx mice treated with vehicle or anti-mIL6 Ab, demonstrating no significant group differences.

|  | Normoxia (Mean±SD)<br>(n=4) | SuHx+Vehicle<br>(Mean±SD) (n=6) | SuHX+anti.mIL6 Ab<br>(Mean±SD) (n=6) |
| --- | --- | --- | --- |
| LV Mass | 0.07±0.008165 | 0.061±0.007528 | 0.058±0.007528 |
| LVEDV | 0.0775±0.012583 | 0.067±0.005164 | 0.072±0.01472 |
| LVESV | 0.125±0.183394 | 0.033±0.005164 | 0.03±0.006325 |
| LVSV | 0.0425±0.009574 | 0.031±0.00482 | 0.043±0.010328 |
| **LVEF** | 56.76±1.399309 | **50.47±4.048224\*** | **58.88±3.44344\*\*** |
| **MAPSE** | 1.178±0.049248 | **0.831±0.134426\*\*** | **1.054±0.147891\*** |

**LV MASS:** Left Ventricular Mass; **LVEDV=** Left Ventricular End-Diastolic Volume; **LVESV=** Left Ventricular End-Systolic Volume; **LVSV=** Left Ventricular Stroke Volume; **LVEF=** Left Ventricular Ejection Fraction; **MAPSE=** Mitral Annular Plane Systolic Excursion

**Supplementary Table 7.**
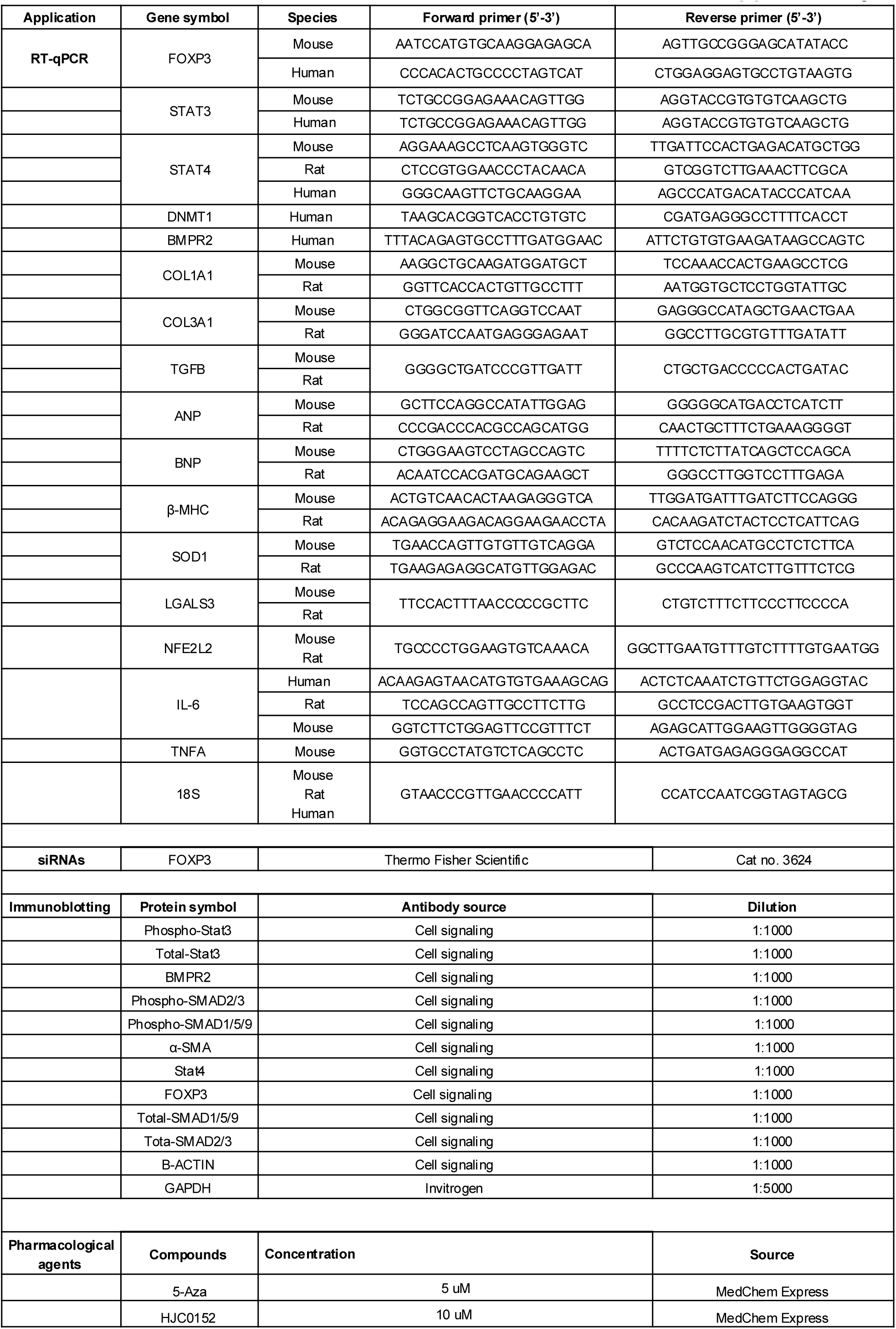
Primer sequences for RT-qPCR analysis, antibodies used, source, and concentration of chemical inhibitors used.

## Supplementary Figures

**Supplementary Figure 1.**
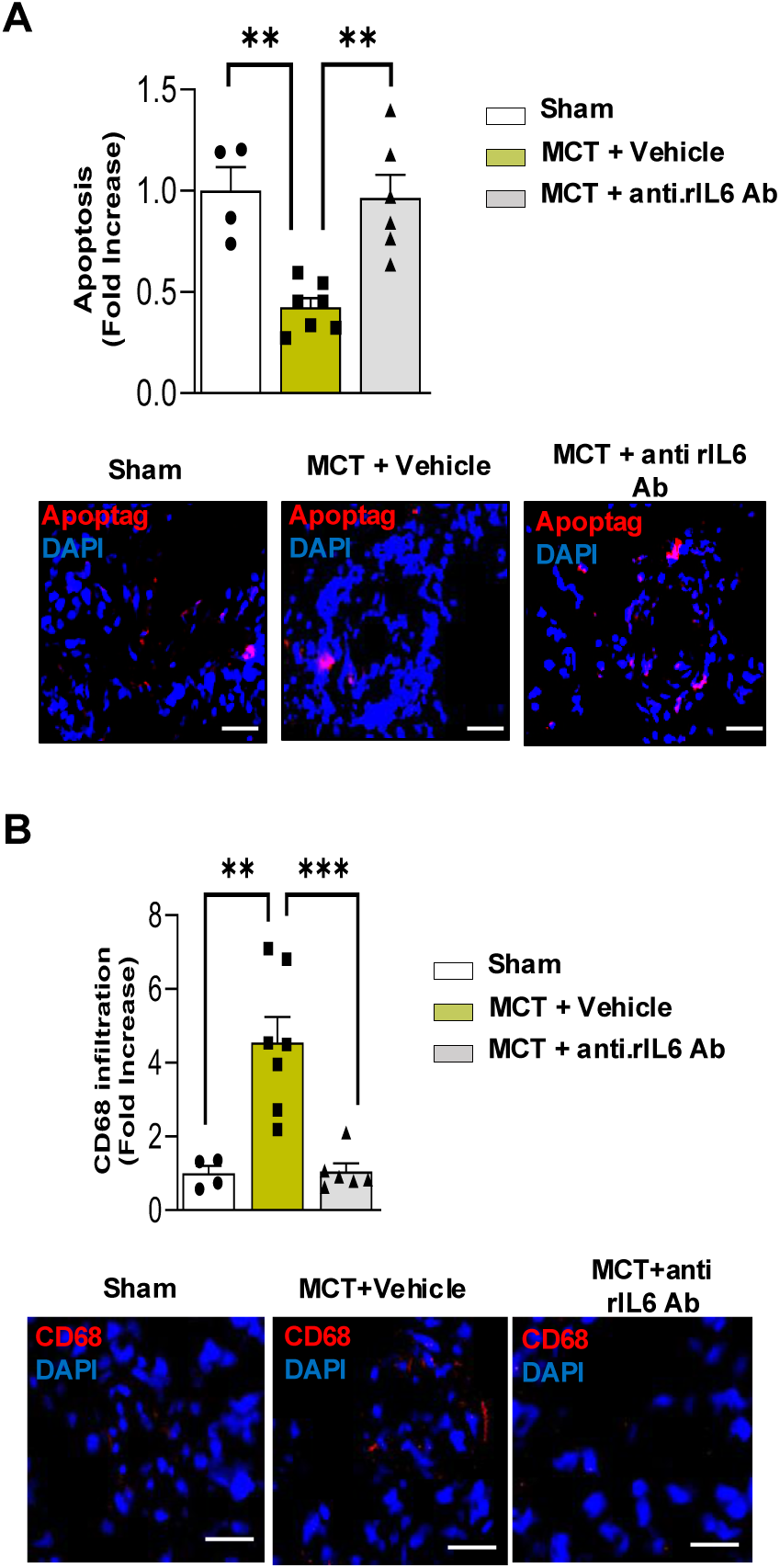
(**A**) TUNEL staining of distal pulmonary arteries demonstrating increased apoptosis in anti-rIL6 Ab-treated MCT rats compared with vehicle-treated controls. (**B**) Representative CD68 immunostaining showing reduced perivascular macrophage accumulation following IL-6 blockade.

**Supplementary Figure 2.**
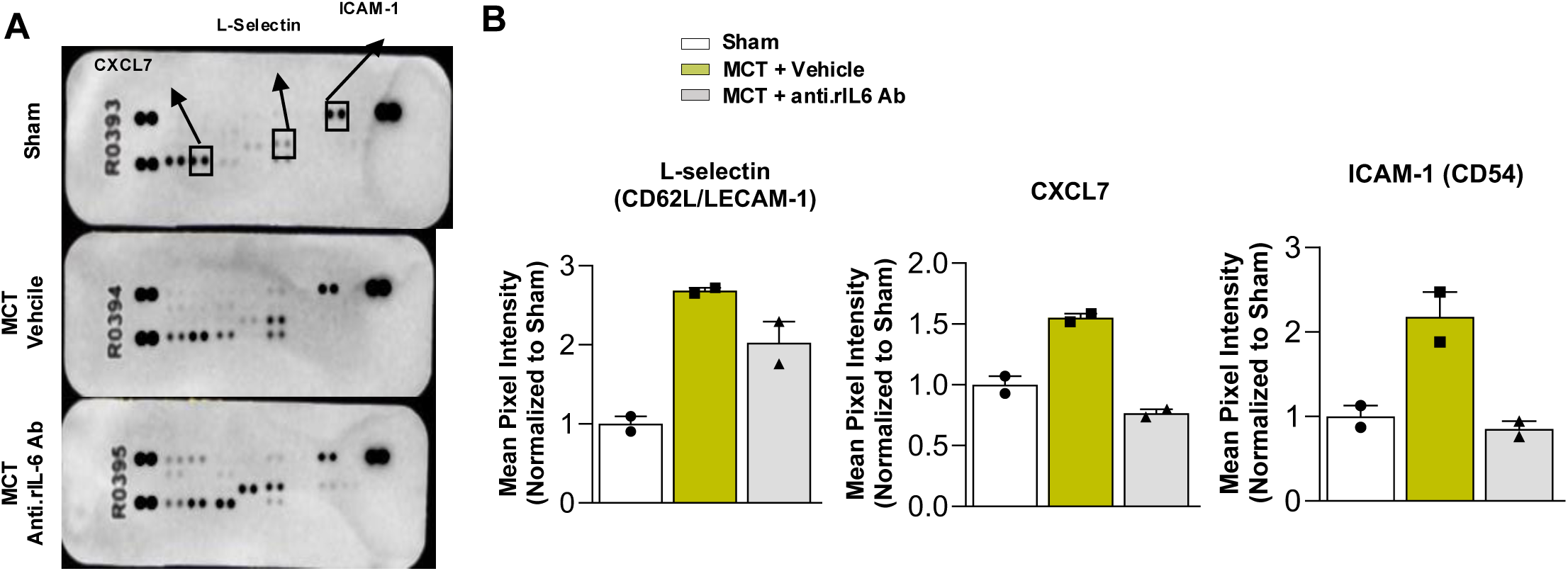
(**A-B**) Cytokine array analysis (Proteome Profiler) of CXCL7 and ICAM-1 in lung homogenates from vehicle- and anti-rIL-6 Ab-treated MCT rats.

**Supplementary Figure 3.**
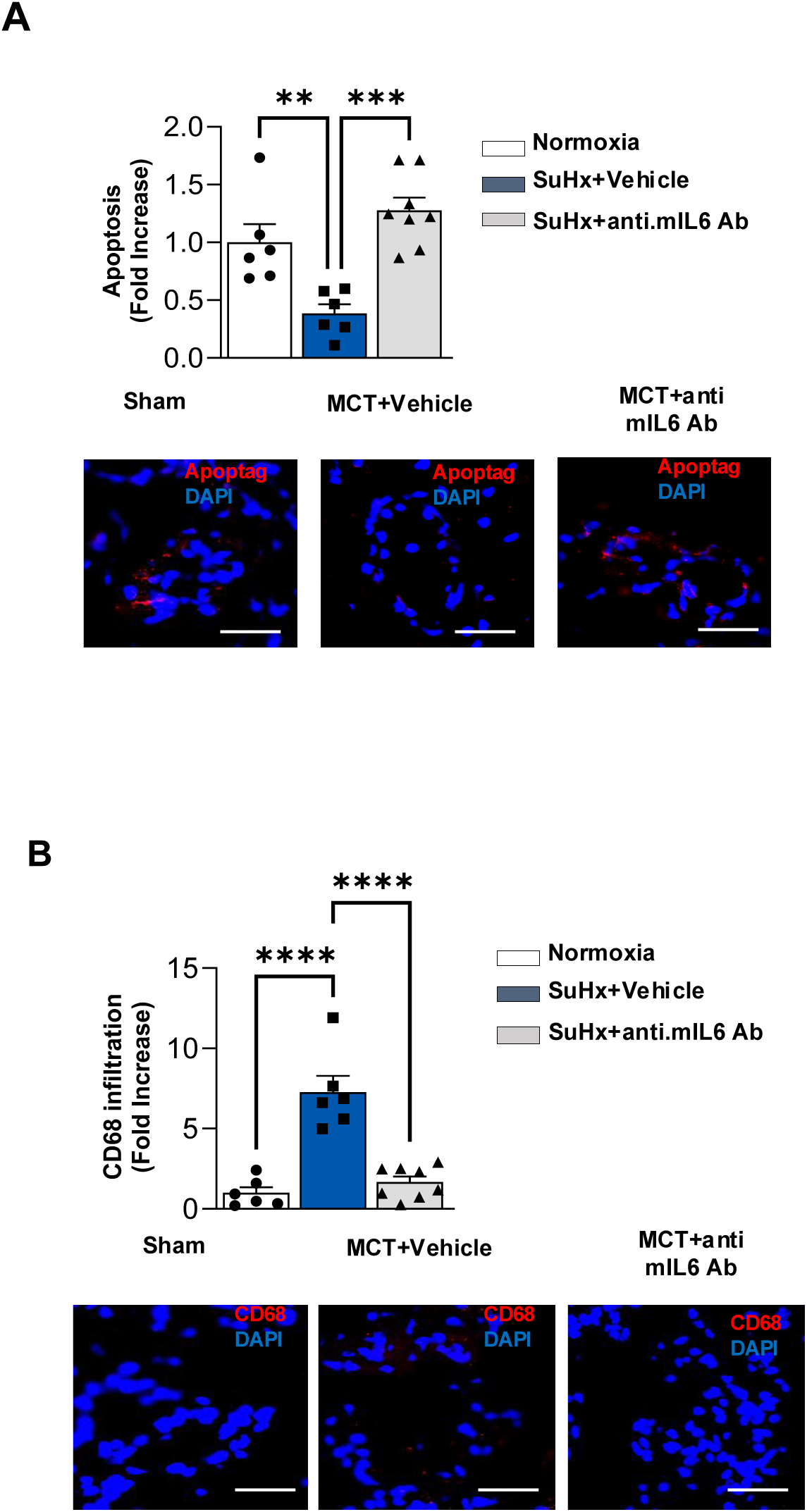
(**A**) TUNEL staining indicates increased apoptosis of hyperproliferative vascular cells following IL-6 neutralization. (**B**) CD68 immunostaining demonstrating reduced macrophage infiltration in lungs of SuHx mice treated with anti-mIL6 Ab.

**Supplementary Figure 4.**
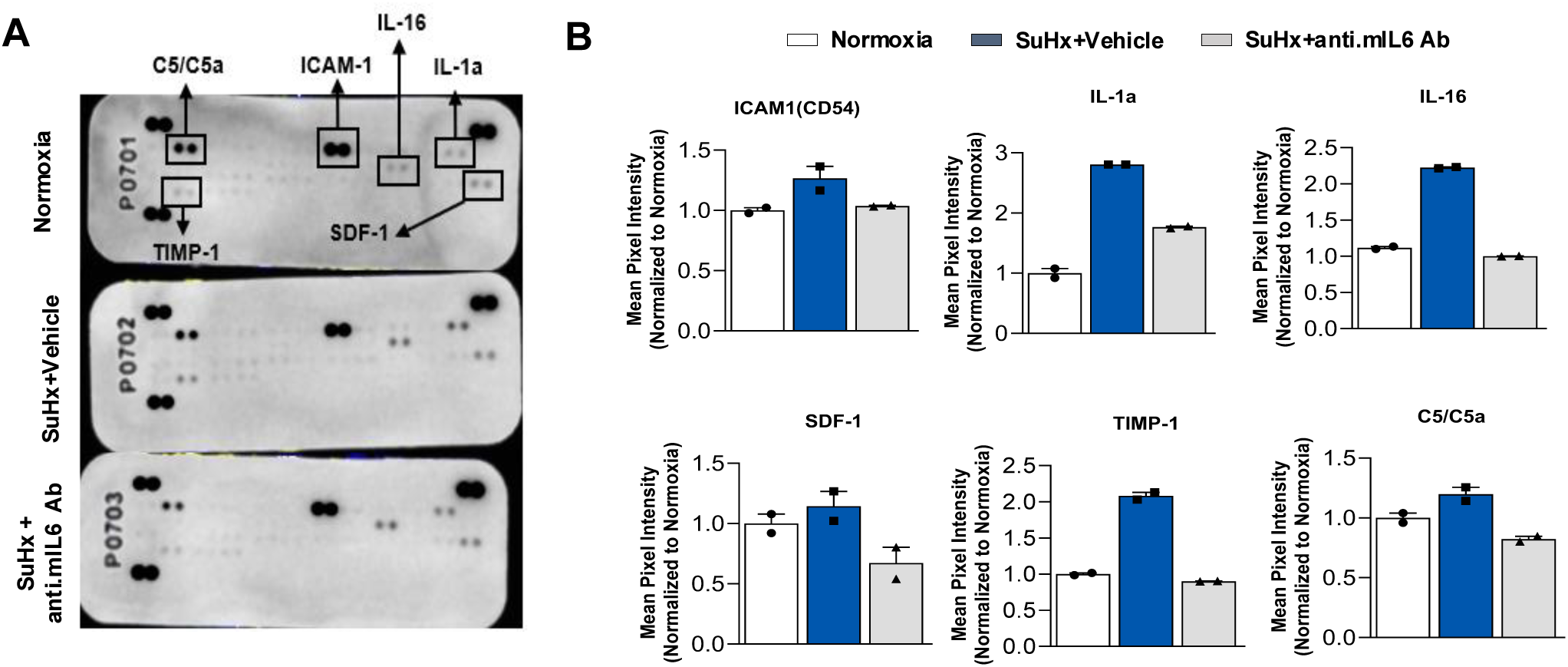
(**A-B**) Cytokine array analysis (Proteome Profiler) of inflammatory chemokines and oxidative stress markers in lung homogenates from vehicle- and anti-mIL-6 Ab-treated SuHx mice.

**Supplementary Figure 5.**
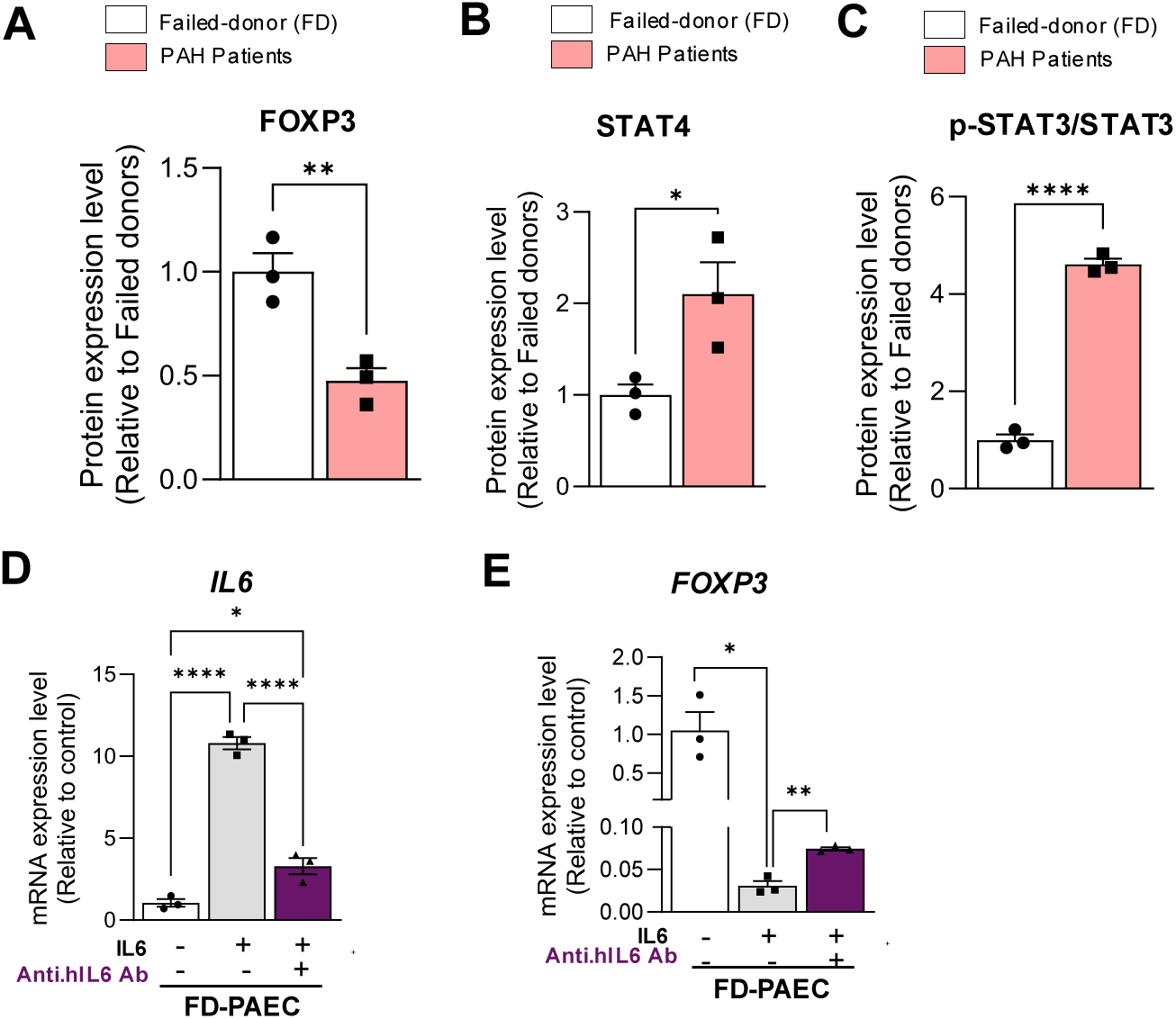
(**A-B**) Densitometric quantification of FOXP3 and STAT4 protein expression normalized to GAPDH in FD-PAECs and PAH-PAECs is shown (n = 3). (**C**) Densitometric quantification of phosphorylated STAT3 (p-STAT3) protein expression, normalized to total STAT3 in FD-PAECs and PAH-PAECs (n = 3). (**D-E**) IL-6 and FOXP3 mRNA expression levels in FD-PAECs treated with recombinant human IL-6, followed by vehicle or anti–hIL6 Ab for 48 hours (n = 3).

**Supplementary Figure 6.**
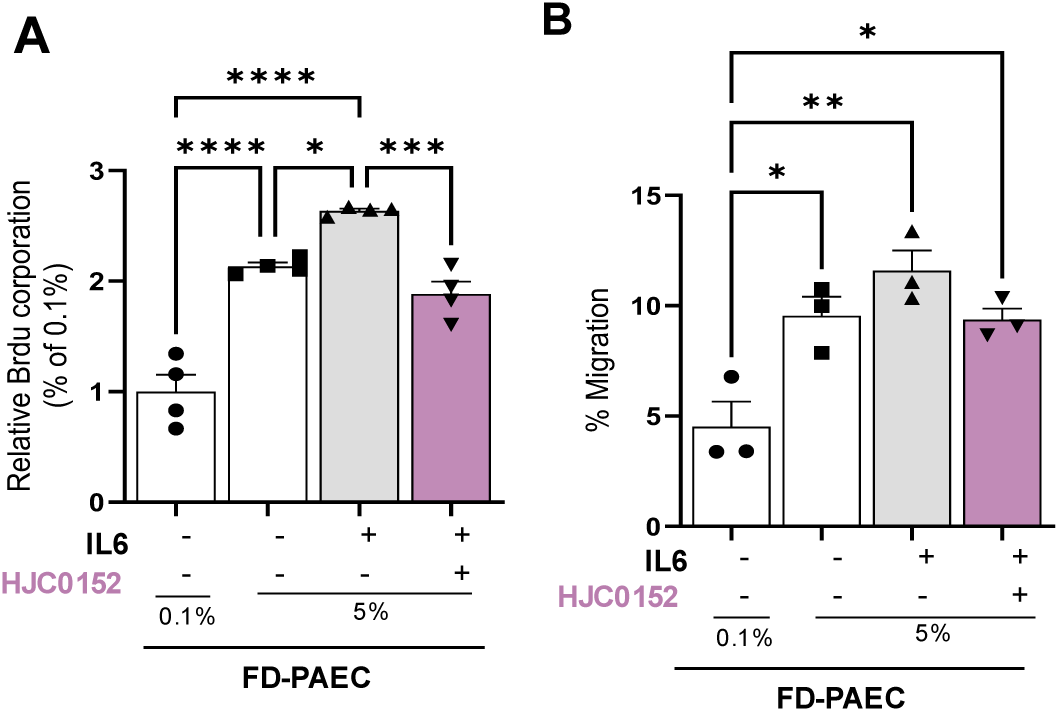
(**A-B**) FD-PAEC proliferation was assessed using the BrDu assay, and migration was assessed using Boyden chambers, respectively. Cells were cultured in high-serum conditions and stimulated with an IL-6 agonist, with or without the STAT3 inhibitor HJC0152, for 48 hours.

**Supplementary Figure 7.**
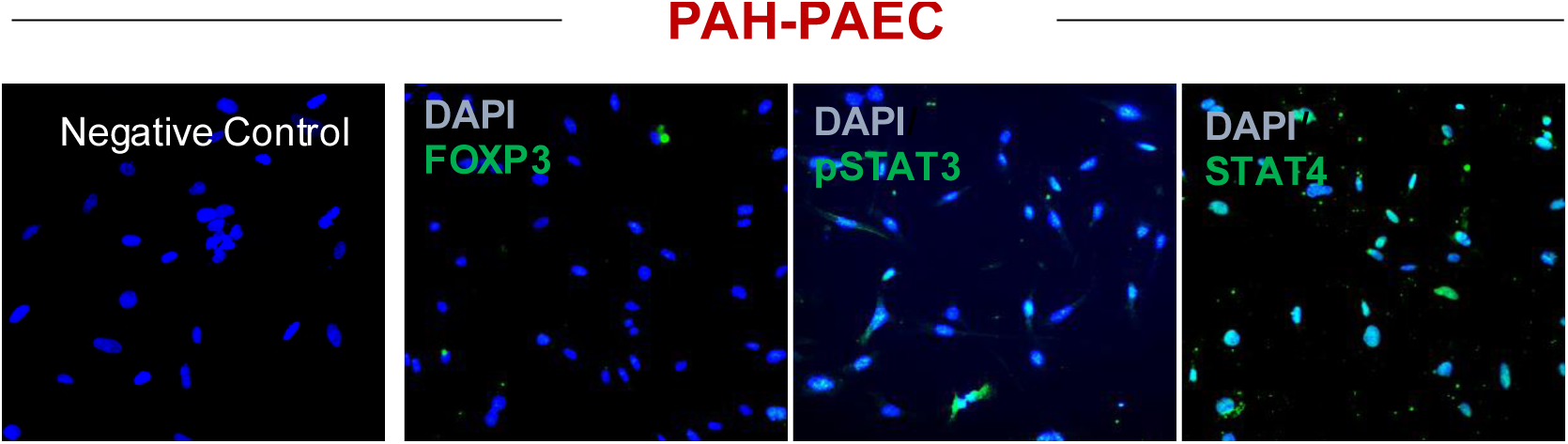
Representative confocal images of immunofluorescence staining for STAT3, STAT4, and FOXP3 (green) in PAH-PAECs. DAPI-stained nuclei are shown (blue). Scale bar = 100 μm.

## List of Abbreviations

Abbreviation: Full Term
α-SMA: Alpha-Smooth Muscle Actin
ANP: Atrial Natriuretic Peptide
BAL: Bronchoalveolar Lavage
BNP: Brain Natriuretic Peptide
βMHC: Beta-Myosin Heavy Chain
cDNA: Complementary DNA
cMRI: Cardiac Magnetic Resonance Imaging
Col1a1: Collagen Type I Alpha 1
Col1a2: Collagen Type I Alpha 2
Col3a1: Collagen Type III Alpha 1
DAPI: 4′,6-Diamidino-2-Phenylindole
ECM: Extracellular Matrix
EF: Ejection Fraction
FOXP3: Forkhead Box P3
H&E: Hematoxylin and Eosin
IL-6: Interleukin-6
IL-6R: Interleukin-6 Receptor
LGALS3: Galectin-3
LV: Left Ventricle
MAPSE: Mitral Annular Plane Systolic Excursion
MCT: Monocrotaline
mPAP: Mean Pulmonary Arterial Pressure
nAb: Neutralizing Antibody
PAEC: Pulmonary Artery Endothelial Cell
PAH: Pulmonary Arterial Hypertension
PASMC: Pulmonary Artery Smooth Muscle Cell
PCA: Principal Component Analysis
RV: Right Ventricle
RVEDV: Right Ventricular End-Diastolic Volume
RVEF: Right Ventricular Ejection Fraction
RVESV: Right Ventricular End-Systolic Volume
RVF: Right Ventricular Failure
RVLS: Right Ventricular Longitudinal Strain
RVSP: Right Ventricular Systolic Pressure
RVSV: Right Ventricular Stroke Volume
STAT3: Signal Transducer and Activator of Transcription 3
SV: Stroke Volume
SuHx: Sugen/Hypoxia
TAPSE: Tricuspid Annular Plane Systolic Excursion
TGF-β1: Transforming Growth Factor Beta 1
Th17: T-helper 17
TNF-α: Tumor Necrosis Factor Alpha
Treg: Regulatory T Cell

## Notes

### Competing Interest Statement

The authors have declared no competing interest.

