## Supplementary material for "Targeting IL-6-STAT3/STAT4 Signaling Restores FOXP3 Expression in Pulmonary Arterial Endothelium and Reveals Novel Biomarkers for PAH": Expanded Methods

To accompany manuscript:

### Expanded Methods

#### Hematoxylin and Eosin (H&E) Staining

The left lung lobe from each animal was fixed in 10% formalin for at least 24 hours, dehydrated, embedded in paraffin, and sectioned at 5 μm using a microtome. Sections were deparaffinized in xylene (3 × 15 min), rehydrated through a graded ethanol series (100%, 95%, 70%), and rinsed in distilled water. Sections were stained with Mayer's hematoxylin for 10 min, rinsed under running water for 5 min, and then incubated with 0.5% eosin for 30 s-1 min. Following staining, sections were dehydrated through sequential ethanol washes (50%, 70%, 95%, 100%), cleared in xylene, and mounted with Cytoseal XYL. Brightfield images were captured with a light microscope, and pulmonary arterial medial thickness was quantified using ImageJ.

#### Immunofluorescence Staining

Paraffin-embedded lung sections were deparaffinized and permeabilized in 0.3% Triton X-100 in PBS for 10 min. Nonspecific binding was blocked with 10% goat serum in Dako antibody diluent for 1 h at room temperature. Sections were incubated overnight at 4°C with primary antibodies, including α-smooth muscle actin (α-SMA, A2547, Sigma) or CD68 (250594, ABBIOTEC). The following day, slides were washed in PBS (3 × 5 min) and incubated with fluorophore-conjugated secondary antibodies for 1h at RT. Nuclei were counterstained with DAPI for 5 min, and slides were mounted using Vectashield antifade medium. Fluorescence images were acquired using a fluorescence microscope.

#### Wheat Germ Agglutinin (WGA) Staining for Cardiomyocyte Size

RV tissue was embedded in optimal cutting temperature (OCT) compound, frozen, and sectioned at 10 μm using a cryostat. Sections were fixed in 4% paraformaldehyde for 10 min and rinsed in PBS (3 × 5 min). Tissue boundaries were outlined using a hydrophobic PAP pen. Sections were blocked for 1 h at room temperature with 10% goat serum in Dako diluent, then incubated with Alexa Fluor 488-conjugated Wheat Germ Agglutinin (WGA, 20 μg/mL) in Dako + 1% goat serum for 1-2 h at room temperature. After PBS washes (3 × 5 min), nuclei were counterstained with DAPI (1:1000 in PBS) for 5-10 min, and slides were mounted with Vectashield. Cardiomyocyte cross-sectional area was quantified using ImageJ software.

**RNA Isolation, cDNA Synthesis, and Quantitative Real-Time PCR (qRT-PCR)**

Total RNA was isolated from the right middle lobe of the lung using TRIzol reagent (Invitrogen) according to the manufacturer's instructions. RNA concentration and purity were assessed using a NanoDrop spectrophotometer. Complementary DNA (cDNA) was synthesized from 1 μg of total RNA using the High-Capacity cDNA Reverse Transcription Kit (Applied Biosystems). Quantitative real-time PCR was performed using Luna® Universal qPCR Master Mix (New England Biolabs) on a real-time PCR system. Gene expression levels were normalized to GAPDH, and relative fold changes were calculated using the ΔΔCt method.

**Western Blot Analysis**

Proteins were extracted from the right superior lobe of the lung using a tissue homogenizer in RIPA buffer supplemented with protease and phosphatase inhibitors. Homogenates were centrifuged at 15,000 × g for 15 min at 4°C, and protein concentration was determined using the Pierce™ BCA Protein Assay Kit (Thermo Fisher Scientific). Equal amounts of protein were separated by SDS-PAGE and transferred onto polyvinylidene fluoride (PVDF) membranes. Membranes were blocked with 5% skim milk in TBS-T and incubated overnight at 4°C with primary antibodies. After washing, membranes were incubated with HRP-conjugated secondary antibodies (Cell Signaling Technology) for 1 h at room temperature. Protein bands were visualized using enhanced chemiluminescence (ECL) reagents (Thermo Fisher Scientific) and quantified using densitometry.

**RNA sequencing and transcriptomic data analysis**

Total RNA was extracted from right lung tissues (n = 3 *per* group) using the RNeasy Mini Kit (Qiagen) according to the manufacturer's instructions. RNA quality and quantity were assessed using a NanoDrop spectrophotometer and an Agilent 2100 Bioanalyzer. Sequencing libraries with 250–300 bp insert sizes were prepared and subjected to paired-end sequencing (150 bp reads) on the Illumina platform. Sequencing was performed by BMKGENE (Durham, NC, USA) using PE150 mode to generate high-quality, comprehensive transcriptomic data. Raw sequencing reads were subjected to quality control and then aligned to the mouse reference genome. Downstream bioinformatic analyses included differential gene expression and pathway enrichment analysis to identify molecular signatures and signaling pathways modulated by IL-6 blockade in PAH.

### Bioinformatics and Data Visualization

Transcriptomic data analysis and visualization were performed using ClusterGrammer (http://amp.pharm.mssm.edu/clustergrammer/), a web-based tool for interactive exploration of high-dimensional data through hierarchically clustered heatmaps. Differentially expressed genes were visualized using volcano plots, with the x-axis representing $\log_2$-fold changes and the y-axis representing $-\log_{10}$-transformed adjusted P-values. Genes with an adjusted P-value $< 0.05$ and an absolute $\log_2$-fold change $\geq 1.5$ were considered significantly differentially expressed. Prior to heatmap generation, raw gene counts were normalized using the logCPM method. The top 2,500 most variably expressed genes were selected after filtering, and a Z-score transformation was applied to standardize expression levels across samples. Hierarchical clustering and heatmap visualization enabled the identification of gene expression patterns and pathway-level effects of IL-6 neutralization in PAH.

### Cell Culture and IL-6 Neutralization

Human pulmonary artery endothelial cells (PAECs) were obtained from PHBI and were isolated from explanted lung tissue or donor lungs (failed donor, FD) not used for transplantation. Briefly, pulmonary arteries were dissected, enzymatically digested, and endothelial cells were purified based on CD31 expression. Cells were cultured in endothelial growth medium under standard conditions and used for in vitro experiments at early passages to preserve phenotypic integrity. FD-PAEC or PAH-PAECs were cultured in endothelial growth medium under standard conditions (37°C, 5% $CO_2$) and used at early passages to maintain phenotypic integrity. Human FD-PAECs and PAH-PAEC were treated with a human IL-6 monoclonal neutralizing antibody (10 µg/mL) or vehicle control for 48 h, depending on the experiment.

### *FOXP3* promoter methylation

FOXP3 promoter methylation was assessed via bisulfite conversion followed by methylation-specific PCR. Genomic DNA was bisulfite-treated, amplified using FOXP3-specific primers, and the percentage of methylation at individual CpG sites was quantified using methylation-specific PCR (MSP) qPCR according to the manufacturer’s instructions. FOXP3 expression and promoter methylation levels were compared between treated and control cells. Correlations with IL-6 exposure and signaling pathway activation were assessed using standard statistical methods.

### Co-immunoprecipitation assays

STAT3-STAT4 interactions were assessed by co-immunoprecipitation using nuclear extracts from PAECs. Equal amounts of nuclear protein were incubated with antibodies against STAT3 or STAT4 overnight at 4°C, followed by capture with protein A/G agarose beads. Immune complexes were washed, eluted, and analyzed by SDS–PAGE and immunoblotting with the indicated antibodies.

### Chromatin immunoprecipitation followed by quantitative PCR (ChIP-qPCR)

ChIP assays were performed using the EZ-Magna ChIP™ A/G Chromatin Immunoprecipitation Kit (Millipore, Cat# 17-10086) according to the manufacturer's instructions. Briefly, human PAECs were cross-linked with 1% formaldehyde for 10 minutes at room temperature to preserve protein–DNA interactions, and the reaction was quenched with 125 mM glycine. Cells were then washed with cold PBS, harvested, and sequentially lysed with the Cell Lysis and Nuclear Lysis buffers provided in the kit. Chromatin was sheared by sonication to an average fragment size of 200-500 bp. For immunoprecipitation, sheared chromatin was incubated overnight at 4°C with rotation using antibodies against STAT4, or normal IgG as a negative control. Protein A/G magnetic beads were then added to capture the antibody–chromatin complexes. Following extensive washing to reduce nonspecific binding, the complexes were eluted, and cross-links were reversed by incubation at 65°C for 4 hours. DNA was purified using spin columns included in the kit. Quantitative PCR was performed using Fast SYBR Green Master Mix (Applied Biosystems, Cat# A46112) with primers targeting the FOXP3 promoter region containing predicted STAT-binding sites (primer sequences are listed in **Supplementary Table 1**). Enrichment was calculated as a percentage of input DNA and normalized to IgG controls to confirm specificity. Relative enrichment between experimental groups was expressed as fold change over control conditions.
